# Structure guided mutagenesis of Henipavirus Receptor Binding Proteins reveals molecular determinants of receptor usage and antibody binding epitopes

**DOI:** 10.1101/2023.11.22.568281

**Authors:** KY Oguntuyo, GD Haas, KD Azarm, CS Stevens, L Brambilla, S Kowdle, VA Avanzato, R Pryce, AN Freiberg, TA Bowden, B Lee

## Abstract

Nipah virus (NiV) is a highly lethal, zoonotic henipavirus (HNV) that causes respiratory and neurological signs and symptoms in humans. Similar to other paramyxoviruses, HNVs mediate entry into host cells through the concerted actions of two surface glycoproteins: a receptor binding protein (RBP) that mediates attachment and a fusion glycoprotein (F) that triggers fusion in an RBP-dependent manner. NiV uses ephrin-B2 (EFNB2) and ephrin-B3 (EFNB3) as entry receptors. Ghana virus (GhV), a novel HNV identified in a Ghanaian bat, use EFNB2 but not EFNB3. In this study, we employ a structure-informed approach to identify receptor interfacing residues and systematically introduce GhV-RBP residues into a NiV-RBP backbone to uncover the molecular determinants of EFNB3 usage. We reveal two regions that severely impair EFNB3 binding by NiV-RBP and EFNB3-mediated entry by NiV pseudotyped viral particles. Further analyses uncovered two point mutations (^NiV^N557S^GhV^ and ^NiV^Y581T^GhV^) pivotal for this phenotype. Moreover, we identify NiV interaction with Y120 of EFNB3 as important for usage of this receptor. Beyond these EFNB3-related findings, we reveal two domains that restrict GhV binding of EFNB2, identify the HNV-head as an immunodominant target for polyclonal and monoclonal antibodies, and describe putative epitopes for GhV and NiV-specific monoclonal antibodies. Cumulatively, the work presented here generates useful reagents and tools that shed insight to residues important for NiV usage of EFNB3, reveals regions critical for GhV binding of EFNB2, and describes putative HNV antibody binding epitopes.

## Introduction

In the 1990s, Hendra virus (HeV) and Nipah virus (NiV) were the zoonotic agents behind highly lethal outbreaks in Australia, Malaysia, and Singapore^1–3^. These two viruses became the prototypic members of the Henipavirus (HNV) genus, a member of the paramyxovirus family, and have caused over 20 human outbreaks with a case fatality rate approaching 60%^4^. Since then, a new Hendra virus genotype (HeV-g2)^5,6^ and several new members have been added to the Henipavirus genus, including Mojiang virus (MojV) from Rattus flavipectus in China, Cedar virus (CedV) from Pteropus bats in Australia, and Ghana virus (GhV) from Eidolon helvum bats in Ghana^7–9^. More recently, Langya virus (LangV), a novel Henipavirus with a presumed Soricidae shrew natural host, was identified as the putative cause of a febrile illness in China between 2019 and 2021^10^. The discovery of novel Henipaviruses, recent outbreaks in China and India, and serological evidence of spillover events in Cameroon lend way to concerns about future outbreaks and highlight the need for a better understanding receptor usage and its role in viral pathogenicity^10,11^.

Henipaviruses encode two surface glycoproteins, the Receptor Binding Protein (RBP) and the Fusion protein (F), that facilitate entry. The tetrameric RBP has a head composed of a 6-bladed beta propeller structure that mediates binding and a stalk that mediates dimerization and tetramerization through cysteine bridges^12,13^. Engagement of the RBP with a receptor induces conformational changes that trigger activation of the trimeric F, which undergoes changes to form a pre-hairpin intermediate that exposes the fusion peptide at the viral membrane distal region and subsequently creates a 6-helix bundle to form a fusion pore.

NiV and HeV have been shown to use Ephrin-B2 (EFNB2) and Ephrin-B3 (EFNB3) while GhV only utilizes EFNB2^14–17^. CedV was recently characterized to use EFNB2, ephrinB1 (EFNB1), and select ephrin-As^18,19^. Meanwhile, receptors have not yet been identified for MojV, LangV, or more recently discovered HNV^7,10,20–23^. Previous groups have leveraged alanine mutagenesis or reciprocally exchanged residues to identify amino acids on both the RBP and receptor side important for receptor usage, particularly for EFNB2 ^24–26^. Notably, some approaches identified HeV-S507 as important for EFNB3 binding^27^, NiV-Q533 as a residue important for EFNB2 and EFNB3 binding^25^, and NiV-QY388-89 as a region important for stabilizing the interaction of HNV-RBP with EFNB2^17^. On the receptor side, structural and functional studies have identified that residues within the GH loop of the ephrin-Bs (EFNBs) snugly fit into pickets lined by HNV residues ^19,24,28–30^. A recent study leveraged this GH loop to generate decoy EFNB2 mutants that bind to HNVs but not cognate Eph receptors^31^.

However, both the RBP and receptor studies primarily focus on HNV engagement with EFNB2. The discovery of HNVs that are unable to use EFNB3, but retain EFNB2 usage, enables us to interrogate the molecular determinants of EFNB3 usage. Here, through systematic structure-informed mutagenesis with GhV as an EFNB3-deficient template, we determine the minimal residues essential to prevent EFNB3 usage by NiV and reveal the distinct accommodations made by NiV to interact with EFNB3. With mutants generated in this study, we also identified regions of GhV that play an important role in EFNB2 usage and identify putative binding epitopes for novel GhV-specific monoclonal antibodies.

## Results

### The head domain of HNVs is a critical determinant of EFNB3 binding

To begin elucidating regions that distinguish NiV and GhV usage of EFNB2 and EFNB3, we generated head-stalk chimeras that exchanged the head domain between two RBPs. Others have shown that transfer of the NiV-head to NDV confers EFNB2 binding and usage to NDV^32,33^. However, we first wanted to assess whether transfer of the GhV-head to the NiV-stalk ablates NiV binding of EFNB3 while retaining EFNB2 binding. Using a flow-cytometry based binding assay with soluble EFNB2 (sEFNB2) or soluble EFNB3 (sEFNB3), we observe that transfer of the GhV head domain with truncations at either the base of the globular head (NiV-GhV-Head187) or the stalk (NiV-GhV-Head146) rendered the chimeras unable to bind EFNB3 while preserving EFNB2 binding. The converse mutations in the GhV-backbone (GhV-NiV-Head161, truncated within the stalk, and GhV-NiV-Head-203, truncated at the base of the globular head) conferred EFNB3 binding while maintaining EFNB2 binding **(Figure 1 and supplemental figure 1)**. As internal controls for the sensitivity of the flow cytometry assay, we included two HeV-RBP constructs that have been previously described to impart differences in EFNB3 binding. Consistent with the literature, we observe statistically significant reduction in EFNB3 binding by HeV-RBP bearing a serine at amino acid position 507 (S507), but equivalent EFNB2 and EFNB3 binding is restored upon the introduction of a threonine at that position (S507T)^27^.

**Figure 1.**
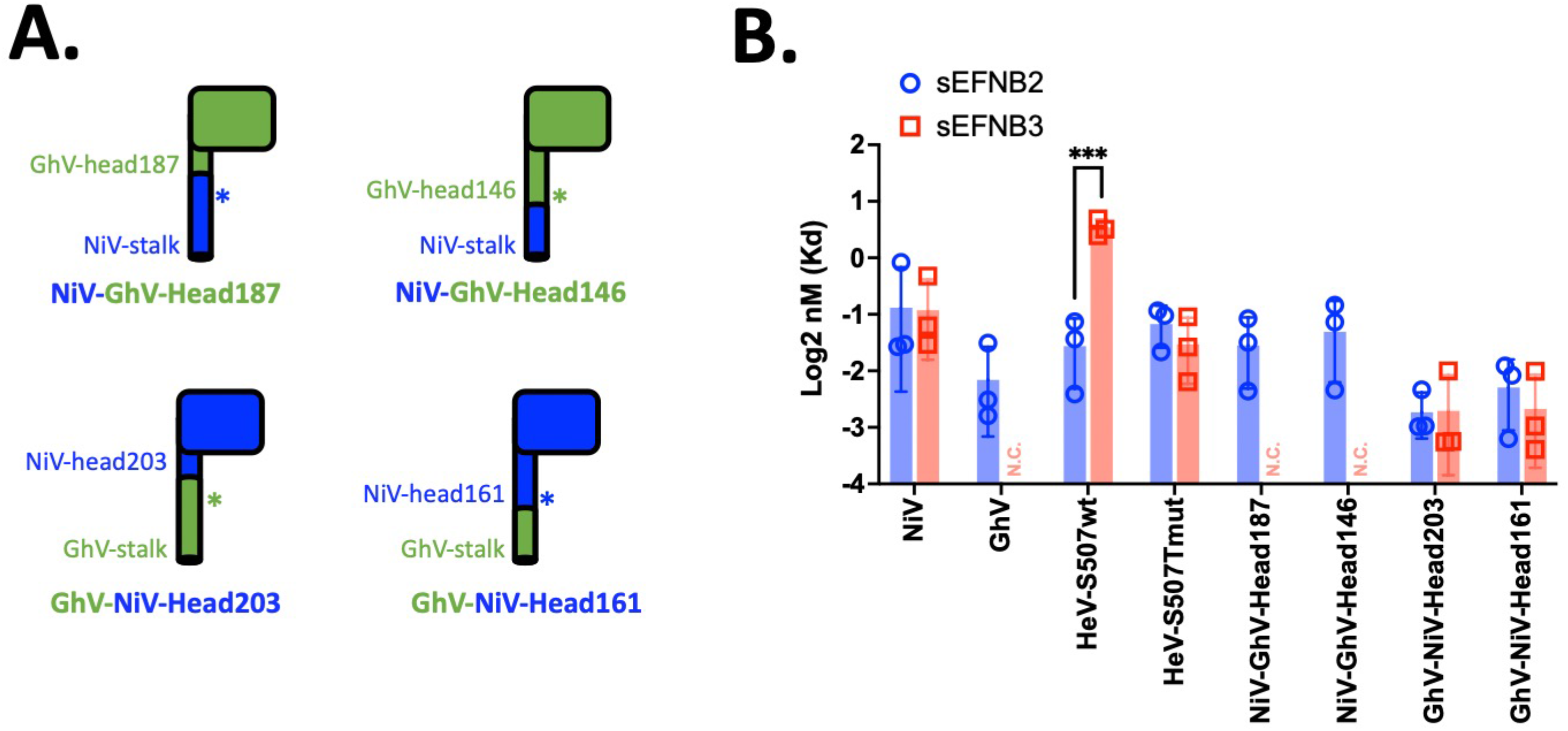
EFNB3 binding localizes to the head domain of HNV RBP. **(A)** Schematic of head-stalk chimeras generated. Constructs presented on top bear a NiV-RBP stalk fused to the GhV head domain at the indicated amino acid position. The two constructs below bear the GhV-RBP stalk fused to the NiV head domain. The asterisks represent the cysteine residues important for HNV tetramerization. Both constructs on the right have a shorter stalk and bear the cysteine residues from the transplanted head. **(B)** Binding of Henipavirus receptor binding proteins to soluble EFNB2 and soluble EFNB3. Flow cytometry was performed by transfecting 293T cells with the indicated HA-tagged HNV RBP then stained with a serial dilution of Fc tagged receptor. This is further described in the Methods. Full binding curves are presented in **supplemental figure 1**. For each point on the curve, the background was first subtracted, then normalized to anti-HA. All data points were further normalized to the highest concentration of soluble receptor tested (50nM). Data were analyzed using a non-linear regression with a saturation binding model with one site. Dissociation constants (kD) presented were calculated from each of three independent biological replicates. N.C. indicates not calculated due to the absence of binding by soluble receptors. An unpaired t-test was performed to calculate statistical significance for all groups with calculated kDs for EFNB2 and EFNB3 (***p &0.0002).

To assess the entry of each chimera into relevant cell lines, we generated VSV-ΔG-RLuc HNV pseudotyped particles (HNVpp) bearing each of the RBPs and homotypic or heterotypic matched F glycoproteins. Consistent with our binding results, we observed the GhV-NiV-Head chimeras were able to enter both ephrin-B2 and ephrin-B3 expressing CHO cells (Cho-B2, Cho-B3). The NiV-GhV-Head chimeras showed no entry into Cho-B3 cells and defective entry into Cho-B2 cells. However, modest levels of entry were observed in the highly permissive U87 cells for 3 of 4 NiV-RBP chimeras and F combinations tested **(Supplemental Figure 2A)**. For the NiV-GhV-head chimeras, we hypothesize that this observed reduced entry may be due to decreased incorporation into the VSVΔG particles **(Supplemental Figure 2B)** or a restricted compatibility of the HNV-RBP chimera stalks with the paired F glycoprotein.

### Systematic, structure-informed mutagenesis reveals residues important for EFNB3 binding and usage

Once established that EFNB3 usage could be ablated in NiV or conferred to GhV by the transfer of the head domain, we leveraged GhV-RBP as an EFNB3-blind henipavirus to further interrogate specific regions necessary to impair EFNB3 usage by NiV. Using a tool to explore molecular interfaces (PDBePISA^34^) with extant crystal structures of NiV, HeV, and GhV RBP in complex with EFNB2^17,28^, we identified residues occluded upon interaction of each RBP with EFNB2. These analyses were also extended to NiV in complex with EFNB3^35^. The results were then visualized on the NiV-EFNB2 structure and annotated onto an alignment of HNV sequences and divided into 12 distinct regions that were subsequently named Occluded Regions (ORs). Interestingly, NiV, HeV, and GhV RPBs share nearly identical interaction profiles for residues occluded upon interaction with EFNB2 or EFNB3 **(Figure 2A)**.

**Figure 2.**
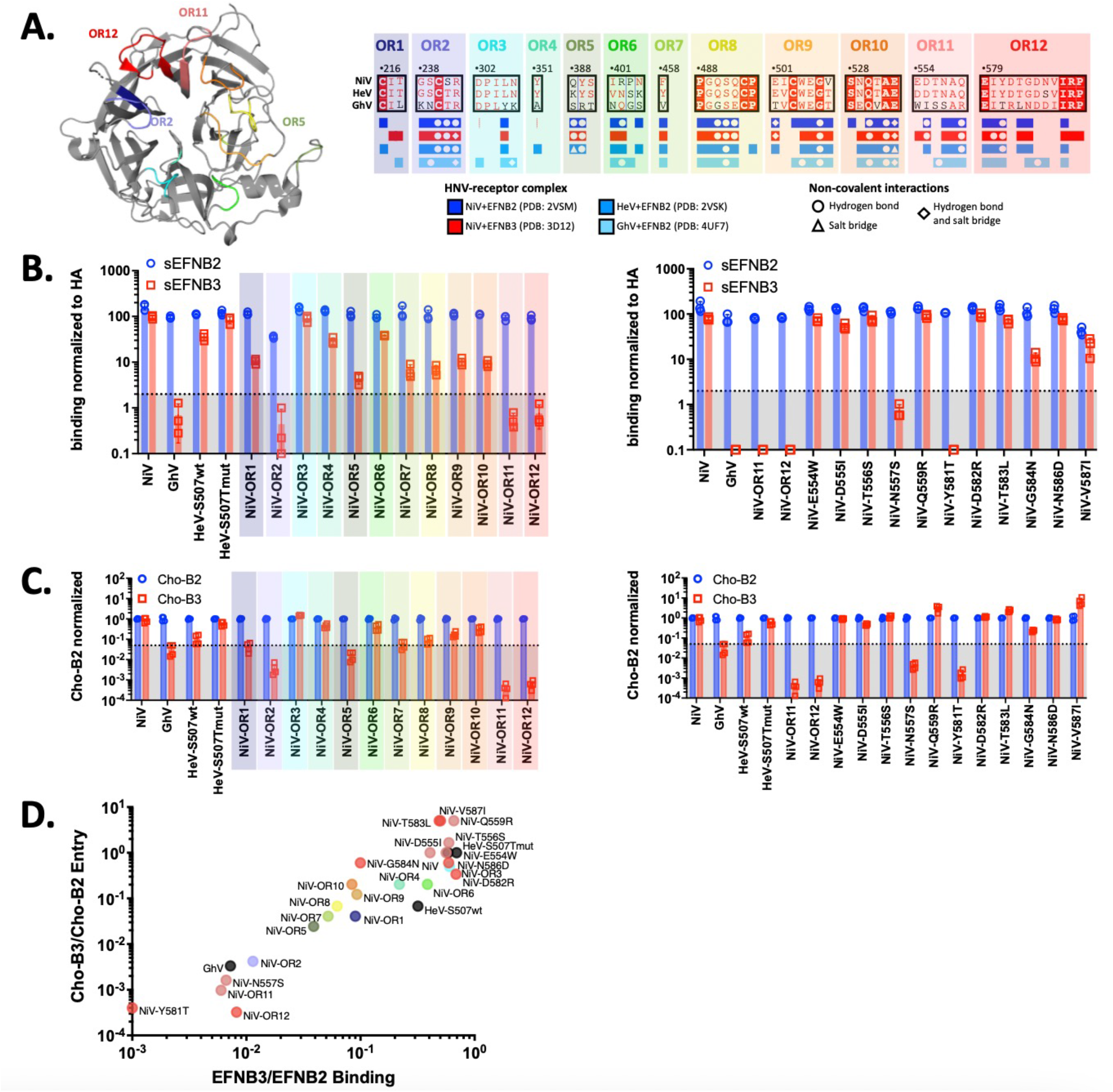
Systematic, structure-guided mutations identify N557 and Y581 as essential for EFNB3 binding and usage. **(A)** Regions occluded by NiV-RBP interaction with EFNB2 displayed on NiV-RBP structure (left) and HNV-RBP alignment (right). Multalin and ESPript were used for multiple sequence alignment and PDBePISA was used to identify occluded regions (OR) 1-12 between the NiV-RBP and EFNB2. Residues occluded in the NiV-RBP and EFNB2 interaction are highlighted in the absence of the EFNB2 receptor for better visualization of their location and regions of interest are noted on the NiV-RBP structure. **(B)** Binding of EFNB2 and EFNB3 by NiV-OR1-12 (left) and selected point mutants (right). Binding was assessed by flow cytometry as described in the Methods. Briefly, 293T cells were transfected with the respective RBPs, then stained with soluble EFNB2, EFNB3, or anti-HA. All RBP are HA-tagged on the external, C-terminal end of the protein, allowing it to serve as a measure of cell surface expression. GMFI from flow cytometry was first background subtracted to account for any non-specific binding, then normalized to anti-HA for each soluble receptor. Since data was first background subtracted, all with a negative value or value &0.01 were given a value of 0.01 to reflect no binding and be presented on the logarithmic graph. These data show binding of soluble receptor at 2nM but binding at 10nM and 0.4nM are shown in **supplemental figure 3**. Presented are the results from three independent biological replicates. The dotted line is at 2 to visualize a stringent threshold for receptor binding. Assuming receptor binding is 100% of HA binding, this indicates an approximately 50x reduction in receptor binding. The shaded region indicates regions that fall below that value. **(C)** Entry into Cho cells stably expressing EFNB2 (Cho-B2) or EFNB3 (Cho-B3) by NiV-OR1-12 (left) and selected point mutants (right). HNVpp bearing the respective control RBPs with homotypic F glycoproteins were prepared using the VSVΔG-rLuc pseudotyping system as described in the Methods. Clarified supernatants containing HNVpps were tittered using a 5-fold serial dilution on Cho-B2 and Cho-B3 cells starting at a 1:4 dilution. The dilution that showed RLU values within the detectable limits of the system for both Cho-B2 and Cho-B3 were selected for normalization to average cho-B2 entry. Presented are the results of two independent biological replicates performed in technical replicates (n=4). The full titers are presented in **supplemental figure 5**. OR regions for **Figures 2B left** and **2C left** were highlighted based on the OR1-12 coloring scheme presented in **Figure 2A right**. Entry for NiV, GhV, HeV wt + mut, NiV-OR11, and NiV-OR12 repeated in figure on right for ease of data interpretation. (D) Binding and Entry ratios for EFNB3 to EFNB2. The ratio of EFNB3:EFNB2 binding at 2nM of soluble protein was calculated after normalization as described in Figure 2B and is presented on the X-axis. NiV-Y581T was given a binding ratio of 0.001 due to having a negative value after subtracting the GMFI background. The ratio of EFNB3:EFNB2 titers was calculated from the average titers on Cho-B2 and Cho-B3 cells presented in **supplemental figure 5**. Each point is colored based on the coloring scheme presented in Figure 2A right and control samples (GhV, NiV, and HeV) have black symbols.

We subsequently introduced the GhV-RBP residues from individual 12 ORs into a NiV backbone and assessed the effect on binding with a flow-cytometry assay. Many regions impair EFNB3 binding, suggesting that several amino acids across the receptor binding pocket support EFNB3 binding. However, OR2, OR5, OR11 and OR12 have the most pronounced effect with approximately 20-100x reductions in EFNB3 binding relative to EFNB2 binding **(Figure 2B, left)**. Interestingly, OR2 also impairs EFNB2 binding. This is likely due to the introduction of an N-linked glycan from GhV’s OR2 into the NiV backbone, which was confirmed by the presence of a higher molecular weight RBP glycoprotein on western blot **(Supplementary Figure 4)**. Given OR11 and OR12 displayed the largest reductions in EFNB3 binding, we subsequently introduced individual point mutations from these regions into the NiV-backbone. Here, we observe severely restricted EFNB3 binding by mutations to N557 and Y581. Since we observe only modest decreases in EFNB3 binding by other point mutations, this finding suggests N557S and Y581T are implicated in the phenotype observed with OR11 and OR12, respectively **(Figure 2B, right)**. Binding was also assessed at 10nM and 0.4nM for all constructs. When binding with excess sEFNB3, NiV-OR11, OR12, N557S, and Y581T still displayed drastic impairments in binding **(Supplemental Figure 3)**.

To assess the functional impact of these regional and point mutations on entry into Cho-B2 and Cho-B3 cells, we generated HNVpp bearing each RBP construct and homotypically matched F glycoproteins. As expected from the literature, NiV showed nearly equivalent entry into both cell lines, HeV-S507T had similar entry into both cell lines, HeV-S507 displayed reduced entry into Cho-B3 cells relative to Cho-B2^27^, and GhV displayed no entry into Cho-B3 cell lines^17^ **(Figure 2C)**. Titers were calculated for these control constructs and our chimeric constructs and data were normalized to Cho-B2 entry for each individual construct **(Supplemental Figure 5 and Supplemental Table 1)**. We observe the greatest reductions in Cho-B3 entry for NiV-OR2, OR5, OR11, and OR12 mutants. Specifically, NiV-OR11 and OR12 display over 3000x and 1500x reductions in EFNB3 mediated entry when normalized to Cho-B2, respectively. Within these regions, point mutants N557S and Y581T display approximately 250x and 500x reductions in EFNB3 mediated entry. Although each point mutant accounts for substantial reductions in EFNB3 usage, these findings suggest that mutations from each region work synergistically to have the greatest impact on EFNB3 usage. Moreover, select point mutations, such as Q559R, T583L, and V587I, modestly enhance Cho-B3 entry approximately 2-4x when normalized to Cho-B2 entry **(Figure 2C, right)**. Interestingly, these constructs also display reduced EFNB3 binding **(Figure 2B, right)** but have significantly reduced titers when compared to wild type NiV **(Supplemental Table 1)**. By taking a ratio of EFNB3 to EFNB2 binding and Cho-B3 to Cho-B2 entry data, we summarize our findings for over 25 constructs. Here, it can be clearly appreciated that NiV-OR11, OR12, N557S, and Y581T have the greatest impact on EFNB3 binding and Cho-B3 entry **(Figure 2D)**.

### NiV occluded regions do not confer EFNB3 usage to GhV and shed insight to GhV EFNB2 usage

We next sought to assess if any of these domains are sufficient to confer EFNB3 binding to GhV-RBP by transferring the ORs most implicated in NiV binding of EFNB3. These regions (OR2, 5, 11, and 12) were mapped onto the structure of GhV **(Figure 3A, right)**. In binding studies, all the chimeric GhV constructs (GhV-OR2, OR5, OR11, and OR12) were still unable to bind EFNB3 at soluble protein concentrations of 2nM **(Figure 3B)** and 10nM **(Supplemental Figure 6A)**. Additionally, we transferred several domains from NiV-RBP to generate GhV-OR+ (bearing NiV ORs 2+5+11+12) or GhV-OR++ (bearing NiV ORs 1+2+5+7+8+11+12), but these chimeric constructs expressed poorly at the cell surface and did not bind either EFNB2 or EFNB3 at 10nM **(Supplemental Figure 6D)**.

**Figure 3.**
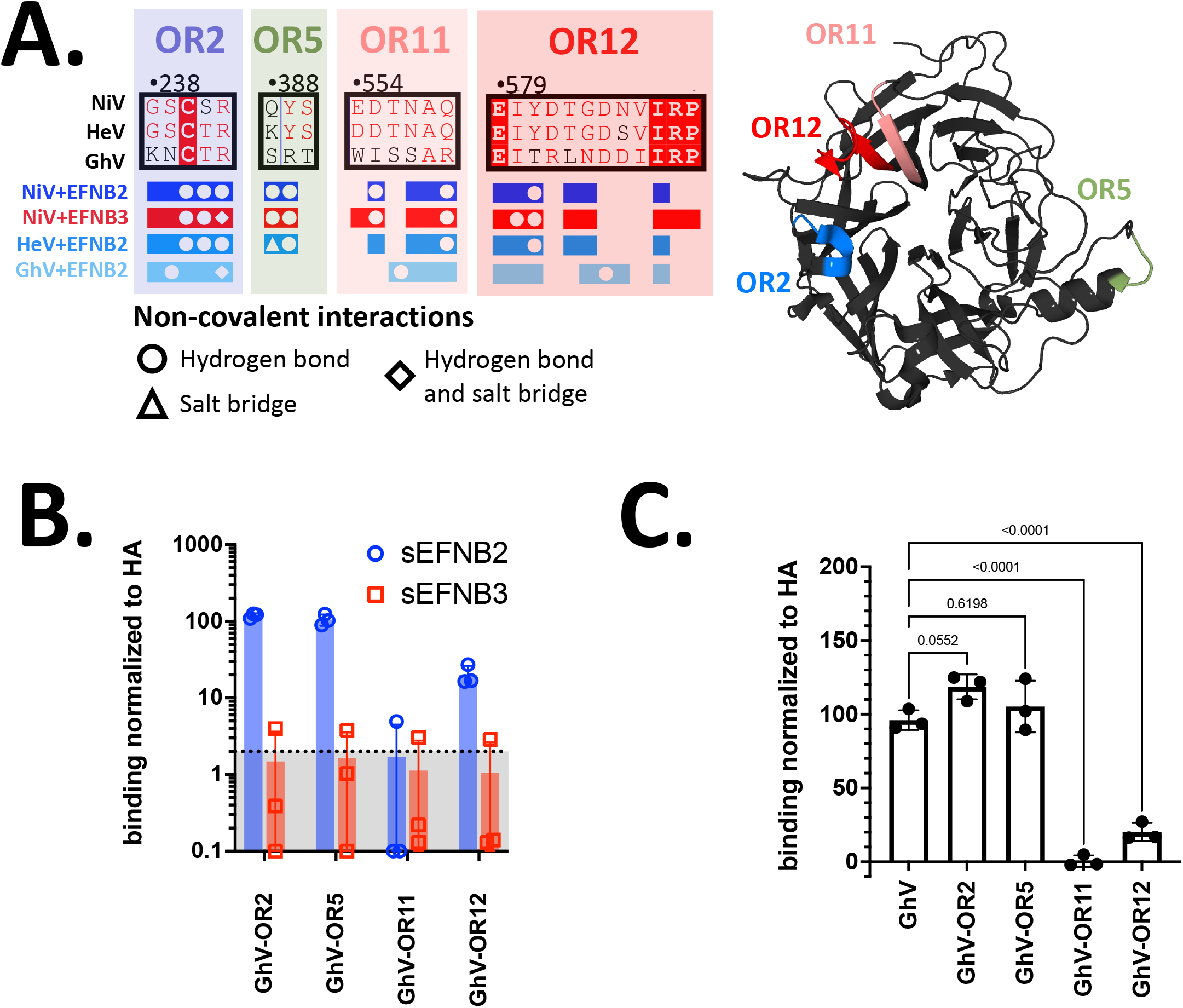
Introduction of NiV RBP residues into a GhV backbone does not confer EFNB3 binding. **(A)** Schematic of occluded regions. Figure 2A was adapted to present occluded regions of interest this figure. Visualization of the localization of each OR region within the GhV-RBP structure. Pymol was used to map the location of OR2, 5, 11, and 12 on the GhV-G structure (PDB: 4UF7). Each region is colored the same as in Figure 2A, **right**. **(B)** Binding of GhV-OR mutants to EFNB2 and EFNB3. Binding of receptors at 2nM is presented here and was performed as described in Figure 2**.3A**. These are the results from three independent, biological replicates. Binding at 10nM and 0.4nM can be found in **Supplemental Figure 6**. **(C)** Binding of GhV-OR mutants to EFNB2. Presented are the data from Figure 3B, but only EFNB2 binding with the addition of GhV-WT-RBP. Statistical significance was assessed with a one-way ANOVA with Dunnet’s correction for multiple comparison. P values are presented atop each comparison.

Further analyses of individual OR mutants in the GhV backbone revealed interesting trends with regards to EFNB2 binding. The removal of the N-linked glycan at OR2 trends towards significant increased EFNB2 binding at 2nM **(Figure 3C)** and has statistically significant EFNB2 binding at 10nM and 0.4nM **(Supplemental Figure 6B)**. Interestingly, GhV-OR11 and GhV-OR12 mutants displayed significant reductions in EFNB2 binding at 2nM **(Figure 3C)** and other concentrations tested **(Supplemental Figure 6B)**. To assess whether this loss of binding is due to disruption of normal GhV conformations, we utilized two RBD-specific, anti-GhV mouse monoclonal antibodies (1E10 and 1G1) to assess cell surface expression of the GhV chimera. Both antibodies bind GhV-OR2, OR11, and OR12. However, these antibodies fail to bind GhV-OR5, suggesting they share an overlapping binding epitope **(Supplemental Figure 6C)**. The structure of GhV-RBP in complex with EFNB2 was reported to have a disordered C-terminal tail that directs upwards towards the receptor binding pocket of the RBP but was not fully resolved^17^. Interestingly, Newcastle Disease Virus (NDV) Ulster strain has a C-terminal extension that is involved in viral entry by mediating dimerization and inhibition of sialic acid binding^36^. Given this and the location of the GhV-RBP disordered C-terminal tail, GhV-OR11 and OR12 mutants may interfere with the orientation of the C-terminal tail and subsequently impair GhV-RBP from maintaining a regional structure necessary for effective receptor binding.

### Accommodation of EFNB3 Y120 is important for EFNB3 binding and usage

Next, we sought to assess why NiV-OR11 and OR12 cannot effectively use EFNB3 while still retaining EFNB2 usage. EFNB2 and EFNB3 are highly conserved across species and in HNV interacting residues **(Supplemental Figure 7)**. However, within HNV interacting residues proximal to OR11 and OR12, EFNB2 bears a phenylalanine at position 117 (F117) and EFNB3 bears a tyrosine at position 120 (Y120). Despite only differing by the addition of a single hydroxyl group found in tyrosine, both residues are buried within a pocket formed at the interface of both OR11 and OR12 **(Figure 4A)**. As a result, we hypothesized that NiV-OR11 and NiV-OR12 chimeras cannot effectively use EFNB3 due to an inability to accommodate Y120. To address this question, we generated cell lines stably expressing wild type (WT) EFNB2, WT EFNB3, or mutant EFNB3 bearing the EFNB2 phenylalanine at position 120 (EFNB3-Y120F). These cell lines were validated as expressing their respective receptor by using flow cytometry to detect binding of EPHB3, a cognate receptor previously reported to bind EFNB2 and EFNB3^37^ **(Figure 4B)**.

**Figure 4.**
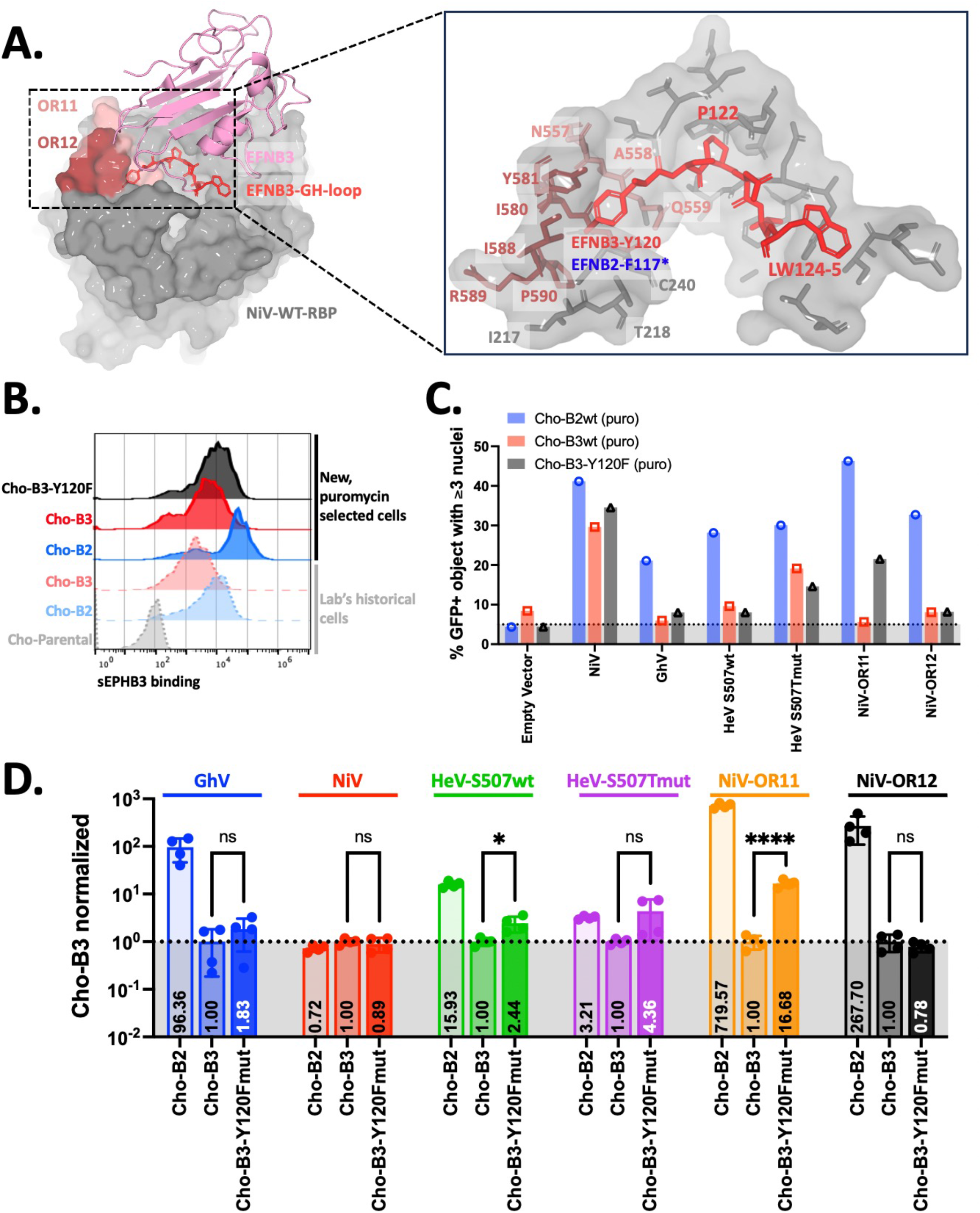
NiV interaction with EFNB3 residue Y120 is important for EFNB3 usage. **(A)** EFNB3- Y120 is primarily enclosed by NiV-OR11 and OR12 residues. Pymol was used to generate complexes of NiV-EFNB3 (PDB: 3D12). EFNB3 in complex with NiV-WT RBP (left) and stick renderings of EFNB3 GH loop residues 120-125 and the NiV binding pocket (right). Asterisk indicates the EFNB2 residue at the same position. **(B)** Cell surface expression of WT EFNB2, WT EFNB3, and mutant EFNB3-Y120F on newly generated cell lines. New, puromycin select cell lines were generated as described in the Methods and compared to historical cell lines for cell surface of each EFNB construct. Cells were stained using a serial dilution of soluble EPHB3 and presented are the histograms from binding to 242nM sEPHB3. The remaining dilutions tested are quantified and presented in **supplemental figure 8. (C)** Quantification of syncytia formed across puromycin selected cell lines. Cells were transfected and imaged as described in **supplemental figure 9**, then put through the CellProfiler pipeline detailed in the Methods. Across each cell line, an average of approximately 500 GFP objects were counted for each transfected well. An object was considered a syncytium if containing ≥3 nuclei within the GFP+ object. The total number of GFP+ objects was then divided by the total number of GFP objects with ≥1 nuclei since a subset of GFP objects lacked a nucleus due to our stringent threshold for including objects on the border of a GFP object. This approach was necessary to avoid overcounting nuclei adjacent to each GFP object. The validation of this system to screen for syncytial phenotypes can be found in **supplemental figure 9. (D)** HNVpp entry into EFNB3-Y120F mutant cell lines. The respective cell lines were infected with the HNVpp and data shown are from the dilutions indicated on the left side. Data were normalized to average Cho-B3 RLUs and presented are the results of two independent biological replicates performed in technical duplicates. Statistics shown are an unpaired t-test (ns = not significant, *p & 0.03, and ****p & 0.0001). Raw RLUs for each biological replicate are presented in supplemental figure 8.

To further validate these cell lines, we developed a GFP based syncytia assay to quantitatively evaluate syncytia formation across several cell lines as a complementary approach to assess for receptor usage prior to producing HNVpp. Briefly, we utilized CellProfiler^38,39^ to count hundreds of GFP+ cells and nuclei within each GFP+ cell per condition tested, then considered syncytia to be GFP+ objects with three or more nuclei. This assay was validated extensively in historical and newly generated Cho-B2 and Cho-B3 cells **(Supplemental Figure 9)**. In the new, puromycin-selected cell lines, NiV displayed similar entry into Cho-B2/Cho-B3 cells and GhV displayed entry only into Cho-B2 cells. Meanwhile, in the puromycin-selected, wild type Cho-B3 cells, GhV, NiV-OR11, and NiV-OR12 showed background levels of syncytia formation. Notably, with NiV-OR11, but not NiV-OR12, syncytia formation was restored in Cho-B3-Y120F cells **(Figure 4C)**.

We subsequently produced HNV pseudotyped particles (HNVpp) and assessed entry into all three puro-selected cell lines. With the newly generated cell lines, control HNVpp continued to display entry in line with previous observations as GhVpp effectively infected Cho-B2 cells and NiVpp effectively infected both Cho-B2 and Cho-B3 cells. Additionally, consistent with our syncytia assay results, NiV-OR11pp, but not NiV-OR12pp, display a statistically significant increase in entry into Cho-B3-Y120F over wild type Cho-B3 cells **(Figure 4D)**. These results emphasize that OR11 plays a role in the accommodation of the hydroxyl-bearing tyrosine residue within EFNB3. Interestingly, when this syncytia assay was expanded to include individual point mutations from OR11 and OR12, we did not observe any Cho-B3-Y120F syncytial phenotype as dramatic as that from NiV-OR11, implicating multiple amino acids as important for this dynamic accommodation **(Supplemental Figure 9F)**. Since NiV-OR12 usage of EFNB3 was not rescued with the Cho-B3- Y120F, we interrogated the structure and identified that Y581T removes a tyrosine from NiV thus preventing pi-stacking necessary to interact with Y120 of EFNB3 **(Figure 5B)**.

**Figure 5.**
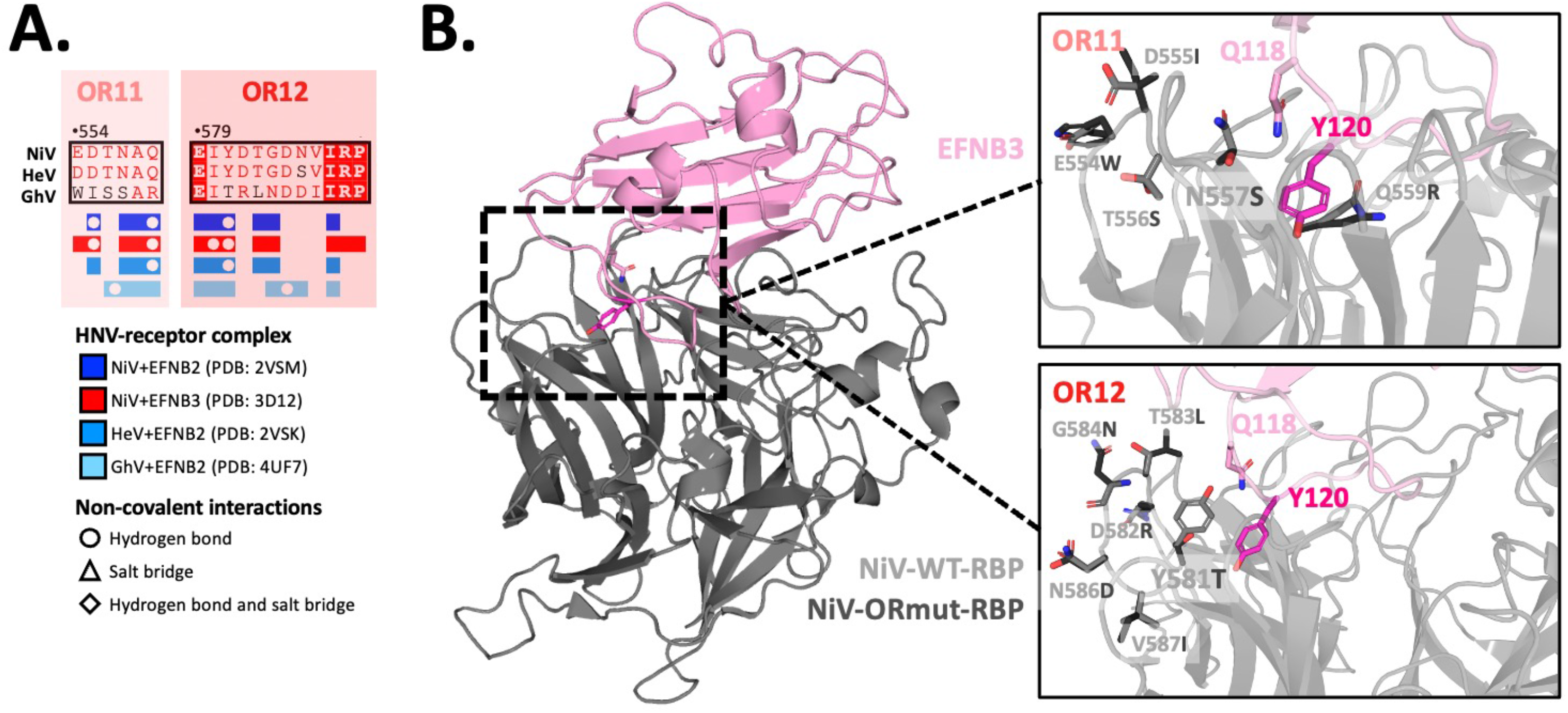
Structural rationale for N557S and Y581T disruption of EFNB3 binding and usage. **(A)** OR11 and OR12 residues occluded by interaction with cognate receptors. This annotation was generated using Mutalin, ESPript, and PDBePISA as described in Figure 2**. (B)** Superimpositions of wild type (WT) NiV-RBP and NiV-OR11 or NiV-OR12 structures. Structures bearing the mutant residues at NiV-OR11, and NiV-OR were generated in COOT. These were then aligned onto NiV-wt-RBP in complex with EFNB3 (PDB: 3D12) in pymol. The full structure is presented on the left with EFNB3 presented in pink and WT-RBP in gray. The right panel shows insets zoomed into a region of the interaction with OR11 (top right) or OR12 (bottom right) with wild type residues presented in light gray and mutant residues presented in dark gray.

### Repurposing head-stalk chimeras and OR mutants to map antibody binding epitopes

Inspired by the findings that OR5 may be the binding epitope for two GhV-RBP specific monoclonal antibodies **(Supplemental Figure 6C)**, we repurposed 20 of the chimeras generated in this study to map epitopes for polyclonal and monoclonal antibodies (pAbs and mAbs, respectively) from our lab. Using the NiV and GhV head-stalk chimeras, we note that all antibodies tested bind to the respective head domains **(Figure 6)**. Although the polyclonal antibodies 2489 and 2491 were generated by immunizing rabbits with DNA encoding full length RBP, none of the antibodies in the serum appear to be stalk reactive. These findings suggest that the head domain of HNVs is immunodominant or the globular head may shield the stalk from recognition by reactive antibodies. Interestingly, NiV pAb806 and NiV pAb2489 display nearly 2- fold increased binding to the GhV-NiV-head chimeras. Similarly, GhV mAb1E10 and GhV mAb 1G1 display nearly 2-fold increased binding to the NiV-GhV-head chimeras. Since these constructs bear different stalk domains, this observation may be due to altered RBP multimerization kinetics.

**Figure 6.**
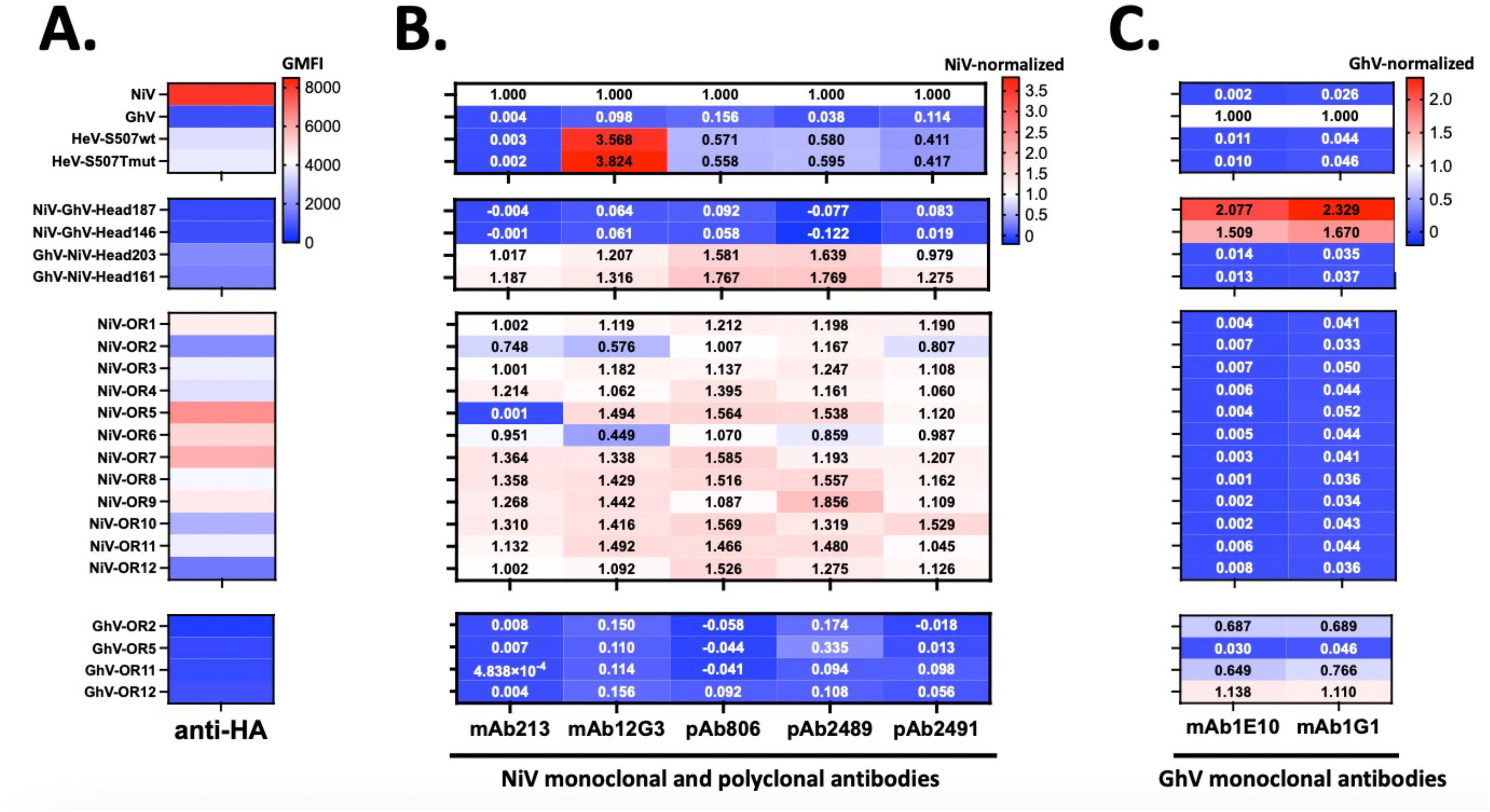
Head-stalk chimeras reveal the immunodominance of HNV-heads and OR mutants identify putative epitopes for HNV antibodies. **(A)** Cell surface expression of constructs as indicated by anti-HA. Flow cytometry was performed as described in the Methods. Briefly, cells were transfected with the indicated HNV RBP, then stained 2 days post transfection. Presented are the background subtracted GMFI from one experiment. **(B)** Binding of constructs to NiV monoclonal and polyclonal antibodies. Flow cytometry was performed as described above and background subtracted GMFI were first normalized to anti-HA GMFI to account for variations in cell surface expression. These values were further normalized to NiV. Presented are the results from one experiment. **(C)** Binding of constructs to GhV monoclonal antibodies. Flow cytometry and analyses were performed as described in panel B of this figure with one exception. For this panel, HA-normalized binding was subsequently normalized to GhV.

We further utilized our NiV and GhV OR mutants to epitope map extant mAbs in our collection as well as GhV specific mAbs that have not been previously characterized. Here, utilizing mAb213 we identify OR5 as a binding epitope, which is consistent with a previous publication that identified Q388R as an escape mutation that developed in live virus escape studies^40^. Additionally, mAb12G3, an antibody originally generated against HeV, shows >3 fold higher binding of HeV compared to NiV. This antibody displays a binding score decreased to approximately 0.5 for NiV-OR2 and NiV-OR6 when normalized to wild type NiV binding. Given this modest reduction in binding and ORs on distinct Beta-sheets, OR2 and OR6 may be epitopes on the periphery of the mAb12G3 binding footprint. Lastly, mAbs 1E10 and 1G1 bind to GhV OR2, OR11, and OR12 mutants. However, both antibodies display decreased binding to GhV-OR5, suggesting this region may represent a binding epitope shared by both antibodies **(Figure 6)**.

## Discussion

Nipah and Hendra viruses have caused outbreaks since the 1990s and manifest clinically with respiratory and neurological signs and symptoms^1^. Of these, the development of neurological signs and symptoms are most strongly associated with mortality^41,42^. Since there is post-mortem evidence of NiV infection of neurons, it is hypothesized that EFNB3, a receptor highly expressed in the central nervous system, may be implicated in the neurotropism and pathogenicity of NiV infections. To better understand the molecular determinants of EFNB3 usage, we leverage NiV (an EFNB2 and EFNB3 using HNV) and GhV (an EFNB2 using, but EFNB3-blind HNV) as HNV-RBPs with distinct receptor usage to characterize regions necessary to impair EFNB3 usage by NiV. Through systematic regional mutations, we identify occluded regions 11 (OR11) and 12 (OR12) as important domains for EFNB3 binding and usage. Through point mutations from these regions, we characterize N557 and Y581 as residues driving this phenotype **(Figure 2).** Moreover, BSL4 experiments with recombinant NiV bearing N557S, Y581T and OR5 mutations continue to display severely impaired EFNB3 usage **(Supplemental Figure 10)** and may prove to be important tools to assess the role of EFNB3 in the neuropathogenicity of NiV.

We further show transfer of single occluded regions did not confer EFNB3 usage to GhV, suggesting that multiple regions or acquisition of non-contiguous mutations may be necessary to enable GhV-RBP binding and usage of EFNB3 **(Figure 3B)**. Interestingly, CedV, an HNV with promiscuous binding to EFNB2, EFNB1, and select ephrin-As, may require fewer mutations to enable EFNB3 usage^18,19^. In fact, Laing et al explicitly highlight that CedV appears to lack the ability to pi-stack with Y120 of EFNB3 due to the presence of N602 in place of Y581 from NiV^18^. This is supported by our data demonstrating Y581T ablates binding and usage of EFNB3, suggesting a shared mechanism by which N602 from CedV and T590 from GhV, both amino acids with polar, neutral side chains, are unable to use EFNB3.

To investigate putative mechanisms for the selective impairment of EFNB3 usage relative to EFNB2, we turned to the receptor side and noted several amino acids buried within binding pockets created by HNV-RBPs. The binding pockets for the LW residues in the GH loop of EFNB2 and EFNB3 are fully formed prior to engagement with the NiV-RBP and have been functionally shown to be critical for usage of EFNB2^16^. However, structural studies suggest that the unbound NiV-RBP appears to make conformational changes to accommodate the tyrosine at amino acid position 120 (Y120) of EFNB3^35^. This residue is buried within a pocket created at the interface of OR11 and OR12. Interestingly, OR12 contains NiV residues 579 to 590, which was previously described in structural studies to represent a dynamic region that undergoes conformational changes upon EFNB2 engagement^43^. Notably, when aligned with EFNB3, EFNB2 bears a phenylalanine (F) at this position so we hypothesized that OR11 and OR12 are unable to bind and use EFNB3 due to an inability to accommodate the hydroxyl group found in tyrosine. We explored the role of EFNB3-Y120 by exchanging this residue with the F found in EFNB2 at this position to generate the mutant EFNB3-Y120F and assessed receptor usage with a novel GFP-based syncytia assay and our standard HNVpp entry assay. Here, in partial support of our hypothesis, we observe that NiV-OR11, but not NiV-OR12, usage of EFNB3-Y120F is partially rescued relative to wild type EFNB3 **(Figure 4)**. While NiV-OR12 usage of EFNB3 is not recovered by the introduction of the Y120F mutation, we hypothesize that it is likely due to an inability for the Y581T mutation to pi-stack with Y120 of EFNB3 consistent with our data with this point mutation and predictions from the literature^18^ **(Figure 5B)**.

From these functional studies, we also made additional observations regarding EFNB2 and EFNB3 receptor usage. For GhV, the introduction of NiV-residues from OR11 and OR12 significantly impaired EFNB2 binding at 10nM, 2nM and 0.4nM despite being cell surface expressed and conformationally intact as indicated by an HA tag and GhV-specific monoclonal antibodies **(Figure 3 and Supplemental Figure 6)**. Though the GhV structure currently lacks resolution of the C-terminal tail, we appreciate the role NDV-Ulster has regulating receptor binding^36^ and hypothesize that OR11 and OR12 play roles to orient the C-terminal tail in a conformation necessary for proper receptor engagement. Notably, truncation of the C-terminal tail has been reported to reduce syncytia formation, suggesting a downstream role in fusion activation^17^.

However, the mechanism driving this phenotype has not been characterized and our results suggest a preceding, putative role of the C-terminal tail in receptor binding. For NiV, we note that NiV point mutants Q559R, T583L, and V587I, show modestly decreased EFNB3 binding, yet display enhanced Cho-B3 entry when normalized to Cho-B2 **(Figure 2 and Supplementary Table 1)**. This discrepancy between binding and entry indicates changes in EFNB3 fusion dynamics with these point mutations. Previous studies have shown subtle conformational changes induced within NiV-RBP upon engagement with EFNB2 through hydrogen deuterium exchange studies and enhanced binding of distinct regions by two monoclonal antibodies^44–46^. While not mechanistically resolved in this work, our results suggest distinct conformational changes induced by NiV binding of EFNB3 compared to EFNB2. This is consistent with our lab’s previous observation that mAb45 binding to NiV-RBP is enhanced in the presence of EFNB2, but not EFNB3^46^. Additionally, a recent structural study highlights a distinct, thumb-like domain that HPIV3-RBP utilizes to engage its fusion glycoprotein prior to fusion triggering and identifies comparable regions within other paramyxovirus that may play similar roles^47^. This region is predicted to be in NiV-RBP amino acids 418-425, which is outside receptor binding pocket for NiV but may be a region that undergoes post-receptor binding conformational changes shared by both EFNB2 and EFNB3.

Lastly, inspired by our fortuitous mapping of putative binding epitopes of two GhV monoclonal antibodies **(Supplemental Figure 6)**, we further leveraged constructs generated in this study to assess binding epitopes for 3 polyclonal antibodies and 4 monoclonal antibodies. Here, we observe that the globular head domain is immunodominant as the NiV-head-stalk chimeras (bearing the NiV-stalk and GhV-head) lose reactivity to the NiV derived polyclonal and monoclonal antibodies **(Figure 6)**. A recent study shows that depletion of head-reactive antibodies from sera from African Green Monkeys immunized with soluble tetrameric NiV-RBP ablates in vitro neutralization potential^48^, which supports our findings that polyclonal antibody binding is lost in the absence of a cognate head domain. Unsurprisingly, introduction of no single occluded region impairs polyclonal antibody binding. Moreover, we identify putative binding epitopes for NiV monoclonal antibodies (mAbs 213 and 12G3) and GhV monoclonal antibodies (mAbs 1E10 and 1G1) **(Figure 6)**. These putative epitopes are supported by the previous report of NiV escape from mAb213 neutralization upon acquisition of a mutation at Q388R, which is within OR5^40^. Epitope mapping is useful for the rational selection of antibodies with distinct epitopes proximal to or outside the receptor binding domain, which is relevant for studies aimed at identifying monoclonal antibody cocktails with synergistic properties for HNV neutralization^48–52^. These constructs also have potential to be utilized as immunogens to generate novel antibodies to distinct NiV regions through use of chimeric OR or point mutant constructs. Moreover, usage of head-stalk chimeras in sequential immunization approaches may lead to the generation of cross-reactive antibodies like vaccination strategies utilized for influenza virus^53,54^.

In sum, the work presented here investigates the molecular determinants of EFNB3 usage, identifying single point mutations that impair NiV usage of EFNB3 and interaction with Y120 of EFNB3 as important for usage of EFNB3. Additionally, we shed insight to regions important for EFNB2 binding by GhV and define antibody epitopes. Importantly, HNV constructs generated in this study may ultimately serve as tools to continue uncovering Henipavirus entry biology, antibody epitopes, and immunogens to generate novel therapeutic antibodies.

## Materials and Methods

### Generation of Receptor Binding Protein (RBP) mutants and EFNB constructs

All constructs were cloned into a pCAGGS backbone utilizing site directed mutagenesis or overlap PCR and were sequence verified prior to use. All wild type fusion glycoproteins were used for these experiments with the exception of the functionally rectified GhV-F previously described^55^. RBP constructs are all codon optimized and contain a C-terminal HA tag that is presented extracellularly. EFNB constructs encode EFNB, a GSG linker, P2A ribosomal skipping sequence and puromycin resistance gene.

### Maintenance and generation of cell lines

HEK293T and U87 cells were maintained in DMEM media. All Cho cells were maintained in DMEM+F12 (Ham’s) media. All media were supplemented with 10% heat inactivated Fetal Bovine Serum. To generate cell lines stably expressing wild type (WT) EFNB2, WT EFNB3, or mutant EFNB3-Y120F, constructs encoding the respective EFNB protein along with a GSG, P2A and puromycin resistance gene were transfected into Cho-Parental cells. These were then put under 10μg/mL of puromycin selection. Cell surface expression of the EFNBs was validated using sEPHB3-hFc (from R&D Cat. No. 5667-B3- 050) prior to freezing down low passage aliquots and using for experiments.

### Structural visualizations, interrogation, and protein modelling

PDB files 2VSM (NiV+EFNB2), 3D12 (NiV+EFNB3), 2VSK (HeV+EFNB2), and 4UF7 (GhV+EFNB2) were utilized for this study^17,28,35^. (Pymol (https://pymol.org) was utilized to generate all the structural models presented here. PDBePISA (https://www.ebi.ac.uk/msd-srv/prot_int/pistart.html)^56^ was used to interrogate the contact residues between HNV-RBPs and cognate receptors. These residues were then annotated onto alignments generated using Multalin (http://multalin.toulouse.inra.fr/multalin/)^57^ and ESPript (https://espript.ibcp.fr/ESPript/ESPript/)^58^. COOT was utilized to mutate select residues as needed prior to visualization in pymol^59^. Superimposed visualizations of structures of interest were generated using the “align” command in pymol given high sequence similarity.

### Cell surface expression and receptor binding by flow cytometry

Cell surface expression of RBP was assessed by transfecting the respective wild type or mutant constructs into HEK293T cells with BioT (Bioland Cat. No. B01-01). Two days post transfection, the cells were gently collected with 10mM EDTA to avoid cleavage of the glycoprotein. Cells were then stained with a 1:2000 dilution of anti-HA (Gentex Cat. No.). For assessing receptor binding, human Fc (hFc) tagged soluble EFNB2 (sEFNB2- hFc) from R&D (Cat. No. 7397-EB-050) or sEFNB3- hFc (Cat. No. 7924-EB-050) was used. Cell surface expression of EFNBs on newly generated cell lines was verified by seeding cells in a 12 well plate prior to collecting with 10nM EDTA and staining with sEPHB3-hFc (R&D Cat. No 5667-B3-050). The Attune was used for all flow cytometry data acquisition and data were analyzed using FlowJo software.

### Production of pseudotyped particles and western blot analyses

Pseudotyped particles were produced as previously described. 293 cells were seeded on poly-lysine coated plates one day prior to transfection with equal amounts of F and RBP or pCAGGS empty vector as a negative control. Eight hours post transfection, these cells were infected with VSVΔG bearing Renilla luciferase for 2hrs and washed three times with DPBS. Media was replenished with DMEM+10%FBS containing a 1:20,000 dilution of an anti-VSV-G antibody 8G5F11 (Kerafast Cat. No EB0010), which reduces any possible background from residual VSV-G. Supernatants were collected and clarified 2 days post infection. In some cases, HNVpp were concentrated through on a 20% sucrose cushion and resuspended in DPBS. Samples were aliquot and stored in -80o to avoid multiple freeze thaw cycles. Western blots were performed on concentrated pseudotyped particles by lysing particles in CA630, resuspending in SDS+BME, boiling the sample for 10mins at 95°C, then running on a 4-15% pre-cast gel. The gel was then transferred to a PVDF membrane and stained with the following primary and secondary antibodies. Membranes were washed with PBS containing 0.1% Tween-20. A mouse anti-VSV-M antibody (Kerafast 23H12 Cat. No. EB0012) was used as a loading control for the particles. Incorporation of HNV-RBP and HNV-F were detected with rabbit anti-HA antibody (Novus Cat. No. NB600-363) and rabbit anti-AU1 antibody (R&D systems nnb600-453), respectively. Goat anti-mouse or anti-rabbit antibodies labelled with Alexa-647 or Alexa-546 was primarily as a secondary antibody.

### Use of CellProfiler for Quantifying Syncytia Assay

CellProfiler (https://cellprofiler.org)^38,39^ was utilized to assess syncytia formation across several constructs in different cell lines. Our lab’s historical Cho-B2 and Cho-B3 cell lines and newly generated Cho-based cell lines were transfected with LifeACT-eGFP (Ref/Cat. No.) and pCAGG empty vector or the indicated HNV F and RBP. This transfection mix contained ∼64% eGFP, 18% F, and 18% RBP. About 22-24hrs post transfection, cells were stained with Hoechst (Abcam ab228551), then fixed with PFA prior to capturing capture brightfield, GFP, and Hoechst images. The Celigo was used to capture the entire well. The same pipeline generated was used to analyze all images in an unbiased fashion. Notably, this pipeline contains a crop function to capture a large field of view (3000×4000 pixels) from the transfected well for analysis. Subsequent steps include identifying primary objects (Hoechst-stained nuclei and GFP+ objects) and masking nuclei to only include nuclei that have at least 50% overlap with GFP+ objects. The latter is a stringent threshold that results in ∼20-30% of GFP+ objects lacking a nucleus since many have nuclei at the periphery of the cells. However, this compromise was made to avoid overcounting nuclei in each GFP+ object. As a result, objects containing no nuclei were excluded from the analyses presented here, particularly when normalizing to the total number of GFP objects. Despite this exclusion an average of >500 GFP+ objects were counted for each condition. After masking nuclei, nuclei were related to the GFP object they were contained within resulting in a “child” (nuclei) to “parent” (GFP+ object) relationship between these two objects. Several measurements were made within the pipeline and exported as a .csv file. Additionally, outlines of GFP objects and nuclei were generated and saved to qualitatively assess the robustness of this system.

## Acknowledgements

We thank all members of the Lee Lab for the conversations that helped guide this project. B.L acknowledges funding from NIH AI123449. K.Y.O. acknowledges funding from NIH F31-AI154739- 01 and NIH T32 AI07647. K.D.A. was supported by NIH F31-FAI133943 and NIH T32 AI07647. G.D.H acknowledges funding from NSF Graduate Research Fellowship and NIH T32 AI07647. C.S. and L.B. were supported by NIH T32 AI07647. T.A.B. acknowledges funding from the Medical Research Council (MR/S007555/). The Wellcome Centre for Human Genetics is supported by Wellcome Centre grant 203141/Z/16Z.

---

**Supplemental Figure 1. Full sEFNB2 and sEFNB3 binding curves for control and Head-Stalk chimeras**. Binding curves from each of the three independent experiments were used to generate individual kDs used in **Figure 1B**. Here, the three biological replicates are aggregated into binding curves for NiV and GhV controls **(A)**, HeV constructs **(B)**, NiV-stalk, GhV-Head chimeras **(C)** and GhV-stalk, NiV-head chimeras **(D)**. For binding experiments, 293T cells were transfected with HA-tagged HNV glycoprotein then stained with a fivefold serial dilution of soluble receptor starting at 50nM and ending at 0.016nM. GMFI from binding were background subtracted, then normalized to anti-HA GMFI. This was further normalized to maximum binding at 50nM. All data points with negative GMFI after background subtraction of maximal binding <0.01 after normalizations were giving a value of 0.01 so that the point, particularly EFNB3 binding, could be visualized on the graph. Graphs show the mean with error bars representing the standard deviation. The dotted lines are from the nonlinear fit and the shaded region represents the 95% confidence bands.

**Supplemental Figure 2. NiV-head confers EFNB3 usage to chimeric GhV-NiV-Head constructs. (A)** Entry of control constructs and homotypically or heterotypically complemented head-stalk chimeras into Cho-B2, Cho-B3 and U87 cells. Homotypically complemented constructs are paired with the stalk-matched F glycoprotein i.e., for NiVwt-RBP a homotypic pairing is NiV-F, and a heterotypic pairing is GhV-F. Cells were infected and processed to read Renilla luciferase generated RLUs as described in the methods. All points shown, except for homotypic NiVwt-RBP, are from a 1:10 dilution of sucrose concentrated HNVpp. Due to high titers in U87 cells, all data for NiVwt is from the 1:10,000 dilution. The dotted line at 500 RLU is indicative of the background we observe with BALDpp (particles produced with Empty vector transfected cells, then infected, and prepared as normal for all other HNVpps) in the Renilla luciferase system. As a result, values not above this threshold are interpreted as no entry in that cell line. Presented are the results from one experiment performed in technical duplicates for Cho-B2 and Cho-B3 cells and technical triplicates for U87 cells. U87 entry was not tested for conditions bearing an asterisk. **(B)** Western blot of incorporation of HNV F and RBP glycoproteins into HNV pseudotyped particles (pp). HNVpp were generated and prepared for western blotting described in the Methods. Anti-HA and anti-AU1 antibodies were used to detect HNV-RBP and F, respectively, then imaged using anti-Rabbit 647 secondary. Anti-VSV-M was utilized as a loading control and detected with an anti-Ms 546 secondary.

**Supplemental Figure 3.** N**i**V **OR regions and point mutants binding at 10nM and 0.4nM sEFNB2 and sEFNB3.** Receptor binding at 10nM **(A)** or 0.4nM **(B)** was performed and analyzed as described in **Figure 2B**. Briefly, constructs were transfected with HA-tagged RBP, then stained with Fc tagged soluble receptor. Data presented are background subtracted GMFI normalized to anti-HA and presented are the results from 3 independent biological replicates. Dotted lines are drawn at 10 **(A)** or 0.4nM **(B)** to visualize a stringent threshold for receptor binding. Bars are colored according to the coloring scheme in **Figure 2A right**.

**Supplemental Figure 4. Incorporation of OR1-12 into HNV pseudotyped particles (HNVpp).** Incorporation of OR1-12 mutants was assessed by western blots as described in **supplemental figure 2A** with one exception. In this western blot, anti-Ms Alexa-647 secondary antibody was used due to the faint signal from the earlier use of Alexa-546 conjugated secondary. Notably, the introduction of GhV residues from OR2 (K**N**CTR) into a NiV backbone (previously bearing GSCSR at that position) introduces an N-linked glycan to NiV-OR2, which is reflected in the modest size increase relative to the other OR mutants. Additionally, GhV residues from OR10 (SEQVAE) remove an N-linked glycan from NiV (previously bearing S**N**QTAE at that position), resulting in a size decrease relative to other OR mutants.

**Supplemental Figure 5. Titering HNVpp bearing NiV-OR mutants and point mutations.** HNVpp were prepared using the VSVΔG pseudotyping system as described in the Methods. HNVpp bearing wt or mutant RBPs and their homotypic Fusion glycoproteins were then utilized to screen for deficits in EFNB3 usage in Cho cells engineered to stably express EFNB2 (Cho-B2) or EFNB3 (Cho-B3). The dotted line at 500 RLUs indicates the background of the Renilla Luciferase assay in our system. Entry not above this threshold is interpreted as no entry. This experiment was done in technical duplicates and repeated once for a biological replicate. Shown are the results of a representative biological replicate with error bars representing the standard deviation of the mean.

**Supplemental Figure 6.** G**h**V **chimeras cell surface expression and receptor binding. (A)** GhV OR2, 5, 11, and 12 binding 10 or 0.4nM EFNB2 and EFNB3. Binding was assessed as previously described in Figure 2.3 and supplemental Figure 2.3. Presented are the results of three independent, biological replicates. Dotted lines are at 10 and 0.4 for binding at 10nM and 0.4nM, respectively. **(B)** GhV binding to 10 or 0.4nM EFNB2. Binding performed and analyzed as described in Figure 2.5D. n = 3 independent biological replicates and p values are indicated on top of each comparison. Notably, OR2 showed no statistical significance at 2nM, but shows statistical significance for EFNB2 binding at 10 and 0.4nM. This suggests GhV-OR2, which removes a glycosylation site from GhV, may improve the binding affinity for EFNB2 and may ultimately improve EFNB2 usage. Additionally, OR11 and OR12 show severely impaired EFNB2 binding even at 10nM. **(C)** Assessing cell surface expression of GhV-OR with GhV-G specific monoclonal antibodies. Two mouse monoclonal antibodies previously generated by Kristopher D. Azarm, PhD in our lab were utilized for binding experiments. Experiments were performed as previously described by transfecting 293T cells, then staining 2 days post transfection with the indicated antibody. Interestingly, OR11 does not bind EFNB2, and appears to be conformationally intact at the cell surface as indicated by these two monoclonal antibodies. However, OR5 binds EFNB2 but is not bound by either antibody tested. **(D)** Cell surface expression of GhV-OR mutants bearing several OR regions from NiV. To assess whether transfer of several domains to the GhV-RBP backbone, would facilitate binding of EFNB3, we constructed GhV-OR+ and OR++ chimeras bearing NiV-OR 2+5+11+12 (OR+) or 1+2+5+7+8+11+12 (OR++). Binding was performed as previously described but presented are the background subtracted GMFI from three independent biological replicates so that the low cell surface expression of GhV-OR+ and GhV-OR++ can be fully appreciated.

**Supplemental Figure 7. Residues occluded on EFNB2 and EFNB3 upon interaction with HNV-RBPs.** PDBePISA, Espript, and Multalin were used as previously described. Several representative animal species were included. Note that this is looking at the residues interacting with HNV-RBPs from the perspective of the EFNB receptors.

**Supplemental Figure 8. Validation of newly generated cell lines. (A)** Cell surface expression of EFNBs on newly generated cell lines. Presented are the background subtracted GMFIs for each of the lab’s historical Cho-B2 and Cho-B3 cell lines and the newly generated, puromycin selected Cho-B2, Cho-B3, and Cho-B3-Y120F cell lines. **(B)** Raw RLUs for NiV-OR11 and OR12 entry into newly generated cell lines. Presented are the RLUs from each infection done in technical duplicates. The dotted line and corresponding shaded region at 500 indicate the background of the Renilla luciferase assay system. As expected, GhV entry into the Cho-B2 and Cho-B3-Y120F cells are at background levels for the system.

**Supplemental Figure 9: Use of and validation of CellProfiler for the automated quantification of HNV syncytia formation. (A)** Schematic of the CellProfiler pipeline. The pipeline was utilized to generate outlines of GFP objects and Hoechst-stained objects. Briefly, the entire 96 well image was loaded, then cropped to be a rectangular object with 3000×4000 pixels for both the GFP and Hoechst image. Mask and relate functions were used to associate “children” nuclei within “parent” GFP objects. More details are available in the Methods. Presented on the left is an image depicting the cropping function for GFP (top left) and Hoechst (bottom left). On the right, is an outline of the GFP object (red outline) and nuclei (green outline). **(B)** Number of nuclei and GFP object area strongly correlate with each other. Here we present the correlation using a simple linear regression between the number of nuclei per GFP+ object and the GFP+ object area. Cells transfected with empty vector and GFP served as a negative control. Control HNV (i.e., NiV, GhV, and HeV) F+RBP and GFP transfected cells served as the positive control. When considering all GFP objects with 1 or more nuclei, 287 objects were counted for Empty vector with an average size of 1238 pixels per object, 420 objects for NiV with an average size of 3408 pixels per object, 656 objects for HeV with an average size of 2905 pixels per object, and 582 objects for GhV with an average size of 1787 pixels per object. A dotted red line and shaded region are at 3 nuclei on the Y axis and 3000 pixels on the X axis to more easily visualize objects falling outside of this region since it contains most of the GFP objects in the empty vector condition. **(C)** Screening all NiV constructs for syncytia formation in historical cho-B2 cell across two technical replicates. Syncytia screening was performed as described above and in the Methods. The number of GFP objects with ≥3 nuclei per syncytia were normalized to the total number of GFP objects with ≥1 nuclei per syncytia. This was performed in technical duplicates to assess the robustness of this assay and hundreds of GFP+ objects were counted per condition. **(D)** Syncytia formation in Cho-B2 and cho-B3 cell lines. To quantitatively assess syncytia formation across cho-B2 and cho-B3 cells the syncytia assay described above was utilized and analyzed as previously. Here, we identified the same trends as previously reported in previous figures using HNVpp. For example, both OR11 and 12 show defects in syncytia formation in cho-B3 cells relative to cho-B2 cells. Presented are the results from one experiment counting hundreds of GFP objects per condition. **(E)** OR11 syncytia formation in puromycin selected stable cell lines. Puromycin selected Cho-B2, B3, and B3-Y120F were transfected with GFP and HNV-F and G glycoproteins. The next day, each well was stained with Hoechst nuclear dye and imaged. CellProfiler generated outlines of each GFP+ object (red) and each nucleus (green). The images presented are a representative segment of the full image that was quantified using CellProfiler in **Figure 4C**. **(F)** Assessment of syncytia formation in newly generated Cho-B2, B3, and B3-Y120 cell lines. The experiment was performed and analyzed as described above. Data are the result of counting hundreds of GFP+ objects per condition. Notably, the new puromycin selected cell lines show the same trends for Cho-B2 and Cho-B3 syncytia formation as the lab’s historical cell lines.

**Supplemental Figure 10. Recombinant NiV with selected mutants display decreased EFNB3 usage in vitro.** Briefly, recombinant Nipah virus (rNiV) bearing the indicated mutants were constructed into a GLuC-P2A-eGFP reporter virus that secretes Gaussia Luciferase. At BSL-4 laboratory facilities, these constructs were rescued on BSRT7 cells using a robust reverse genetics system^60^ and amplified on Cho-B2 cells prior to freezing virus stock at -80. rNiV stock was diluted to from neat to 1/3125, then serial dilutions were used to infect Cho-B2 and Cho-B3 cells prior to collecting supernatants at 24 and 48 hours post infection (hpi). Data from the 1/625 dilution is presented above and was normalized to the Cho-B2 average for the respective sample. NiV-OR5 and NiV-N557S were performed in technical triplicates while NiV-Y581T was performed in technical duplicates.

**Supplemental Table 1. Titers of HNVpp bearing NiV-OR mutants and point mutations.** HNVpp were prepared using the VSVΔG pseudotyping system as described in the Methods and tittered on Cho-B2 or Cho-B3 cells. This was performed in technical duplicates and repeated once for two biological replicates. The table displays the average titer from each biological replicate and a ratio of the ChoB3:ChoB2 titers.

## Notes

### Competing Interest Statement

The authors have declared no competing interest.

