## Supplemental Figure for "Structure guided mutagenesis of Henipavirus Receptor Binding Proteins reveals molecular determinants of receptor usage and antibody binding epitopes"

**Supplemental Figure 1. Full sEFNB2 and sEFNB3 binding curves for control and Head-Stalk chimeras**

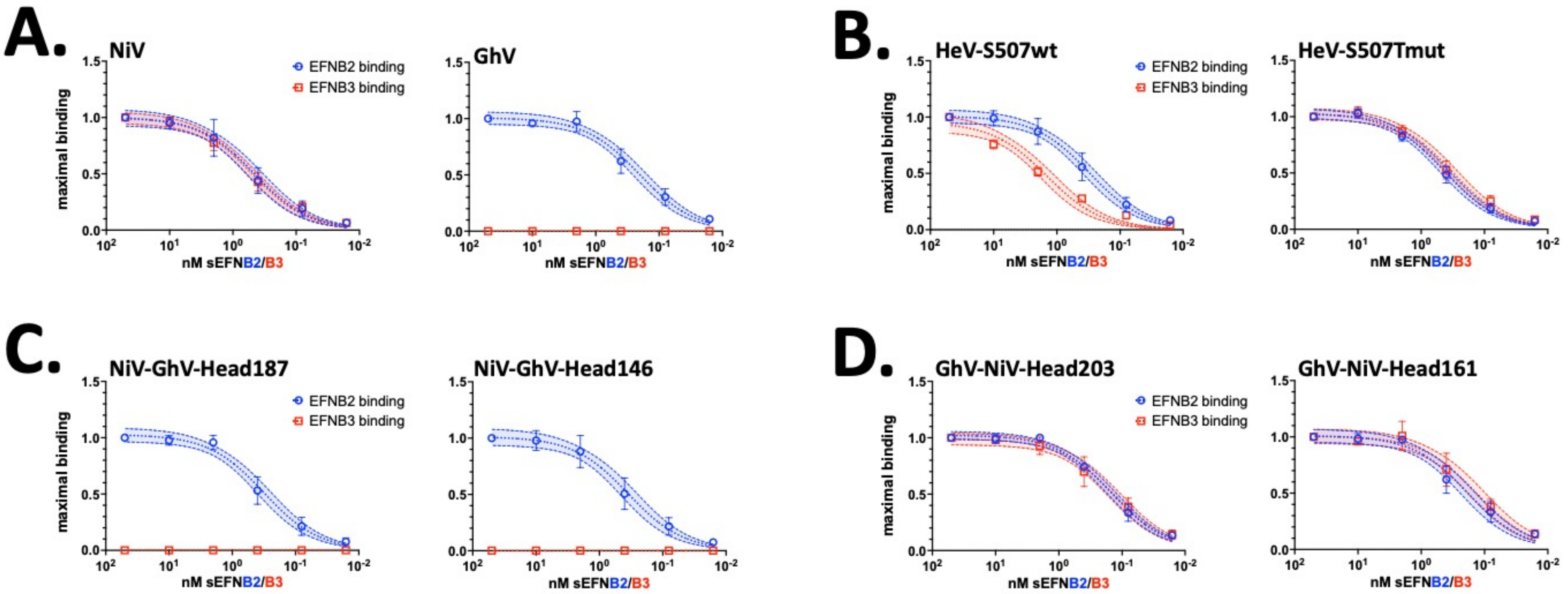

#### Supplemental Figure 2. NiV-head confers EFNB3 usage to chimeric GhV-NiV-Head constructs

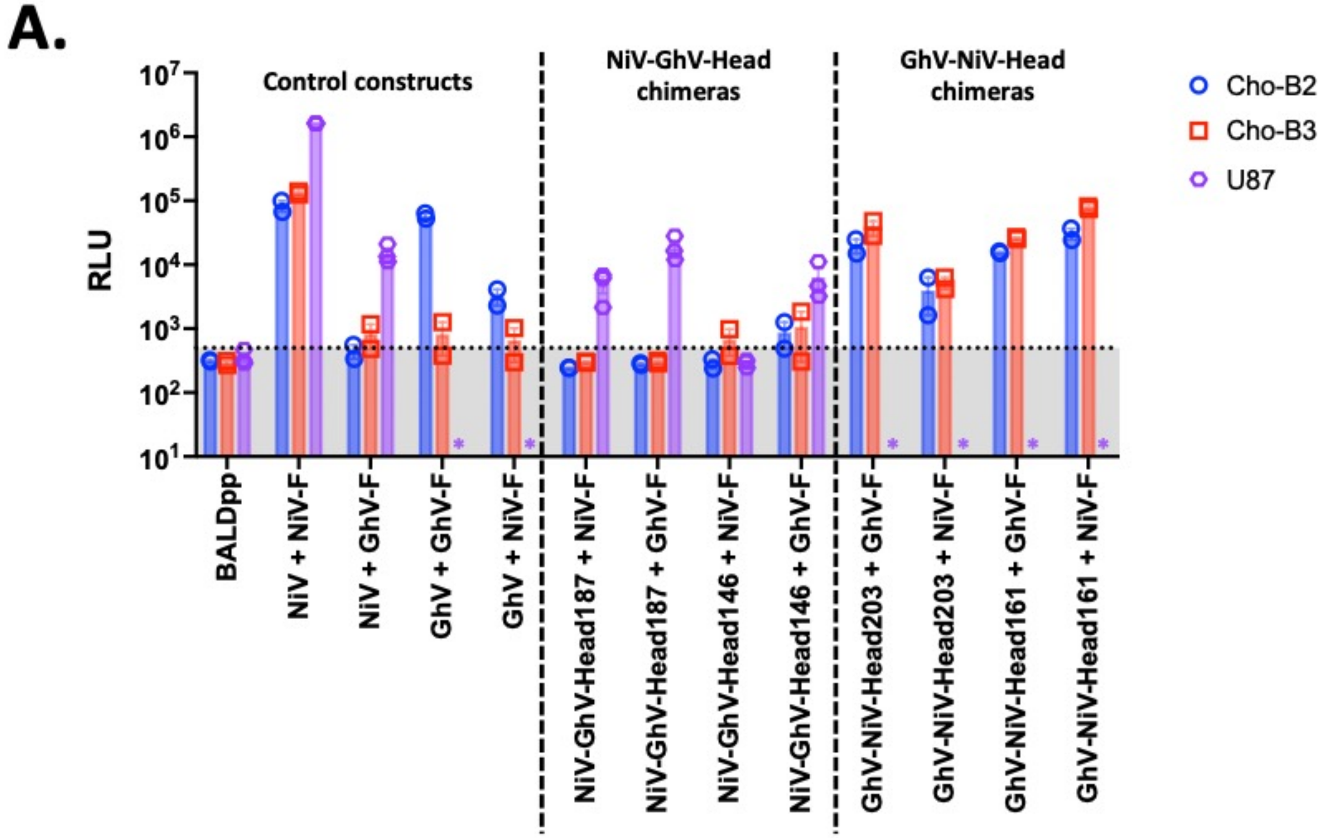


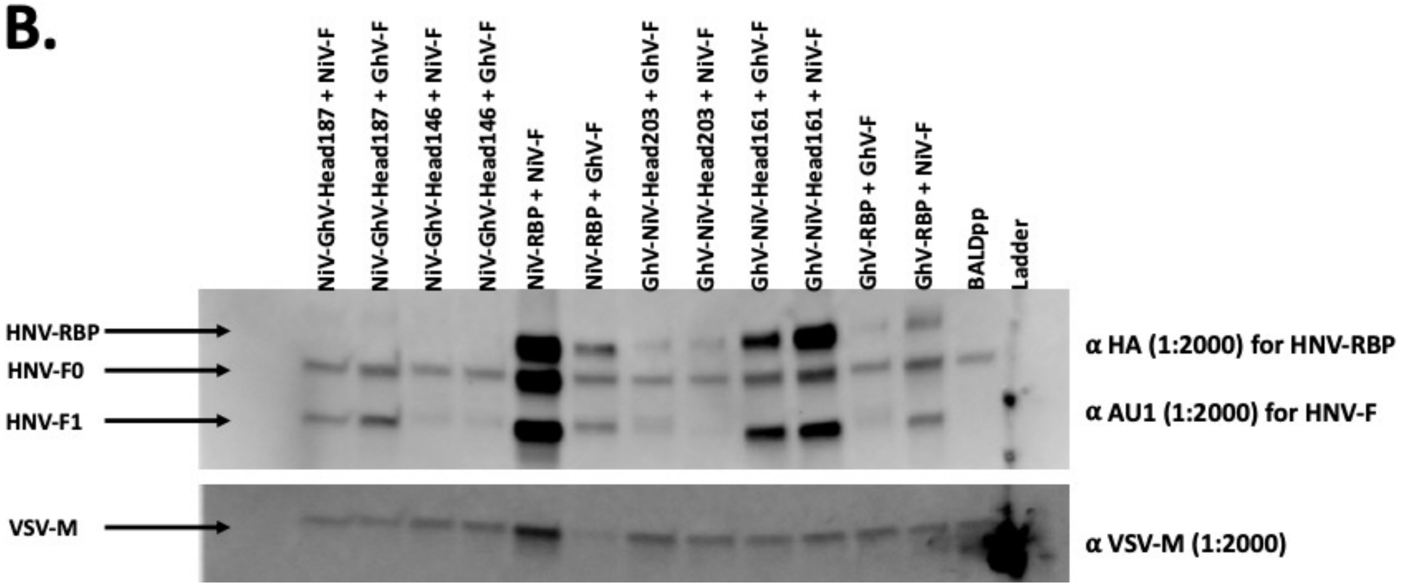

**Supplemental Figure 3. NiV OR regions and point mutants binding at 10nM and 0.4nM sEFNB2 and sEFNB3.**

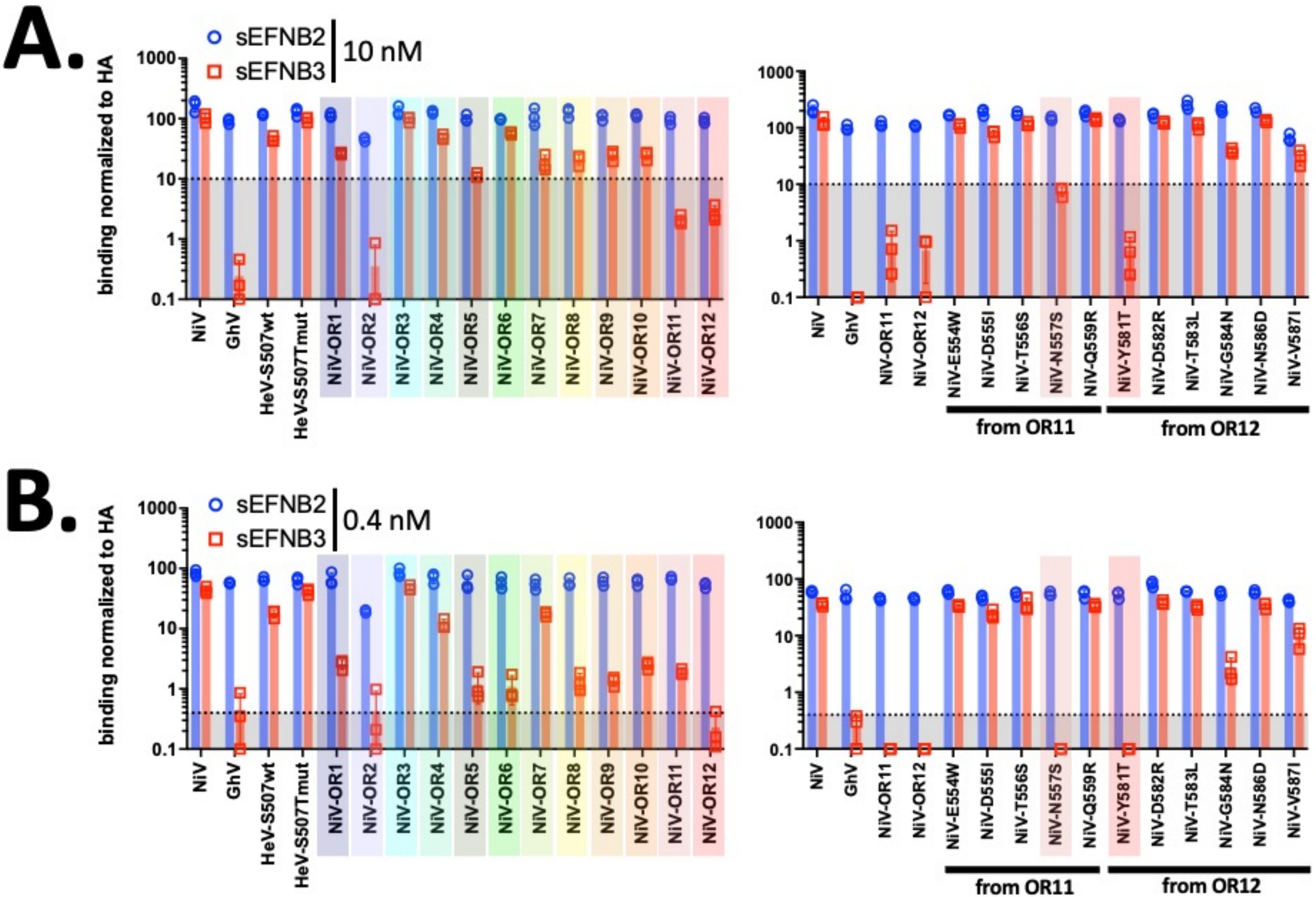

**Supplemental Figure 4. Incorporation of OR1-12 into HNV pseudotyped particles (HNVpp)**

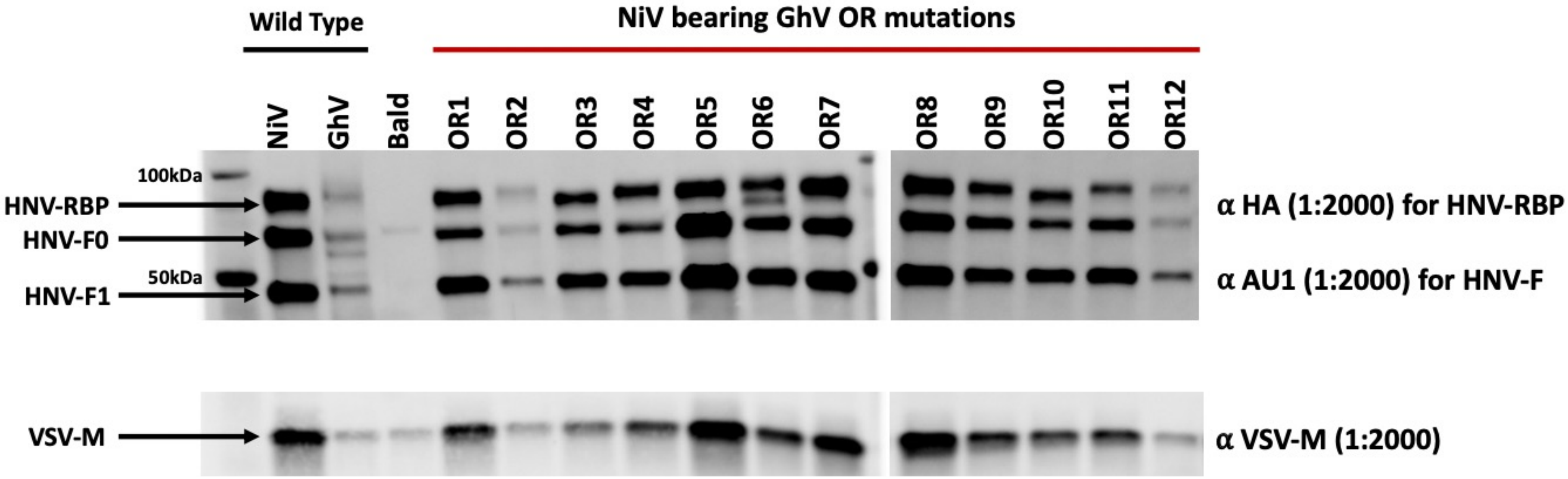

**Supplemental Figure 5. Tittering HNVpp bearing NiV-OR mutants and point mutations**

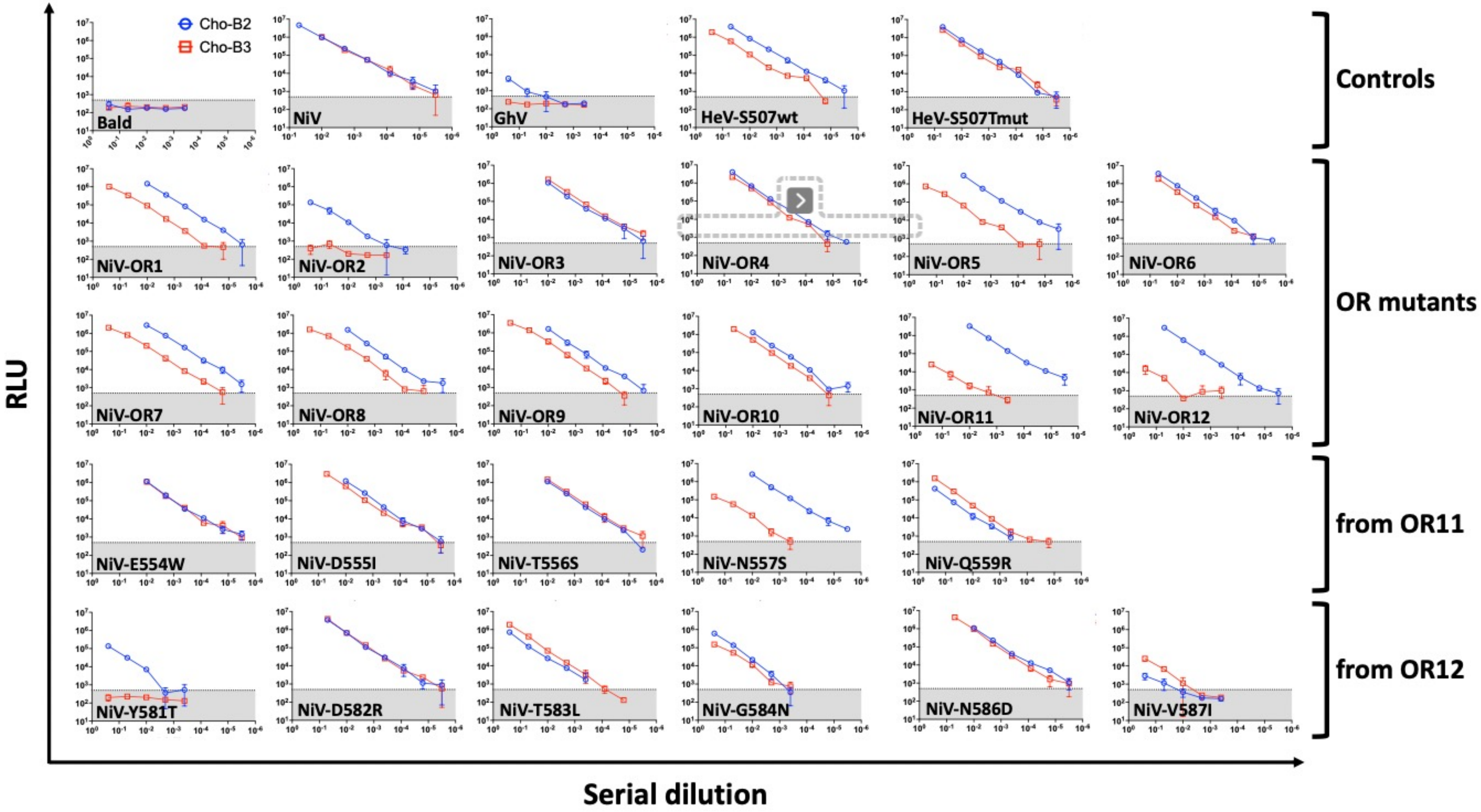

**Supplemental Figure 6. GhV chimeras cell surface expression and receptor binding.**

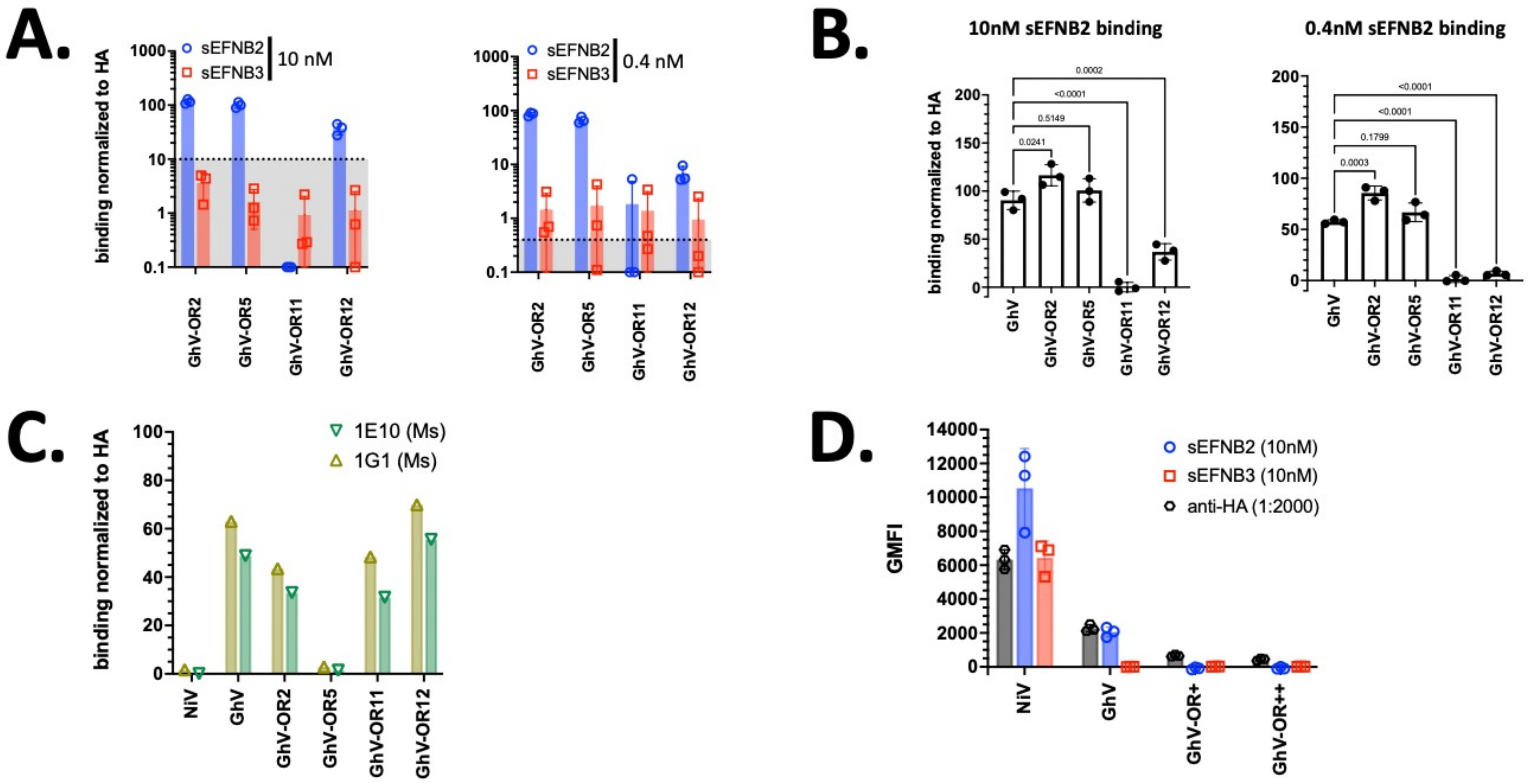

### Supplemental Figure 7. Residues occluded on EFNB2 and EFNB3 upon interaction with HNV-RBPs

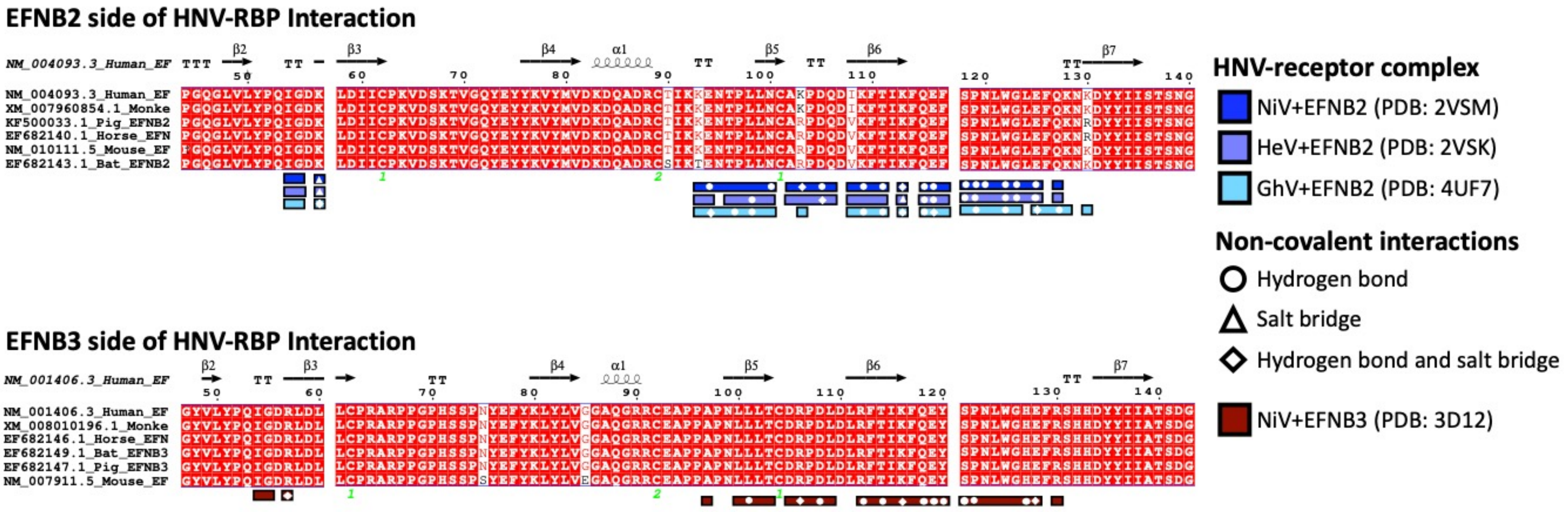

**Supplemental Figure 8. Validation of newly generated cell lines**

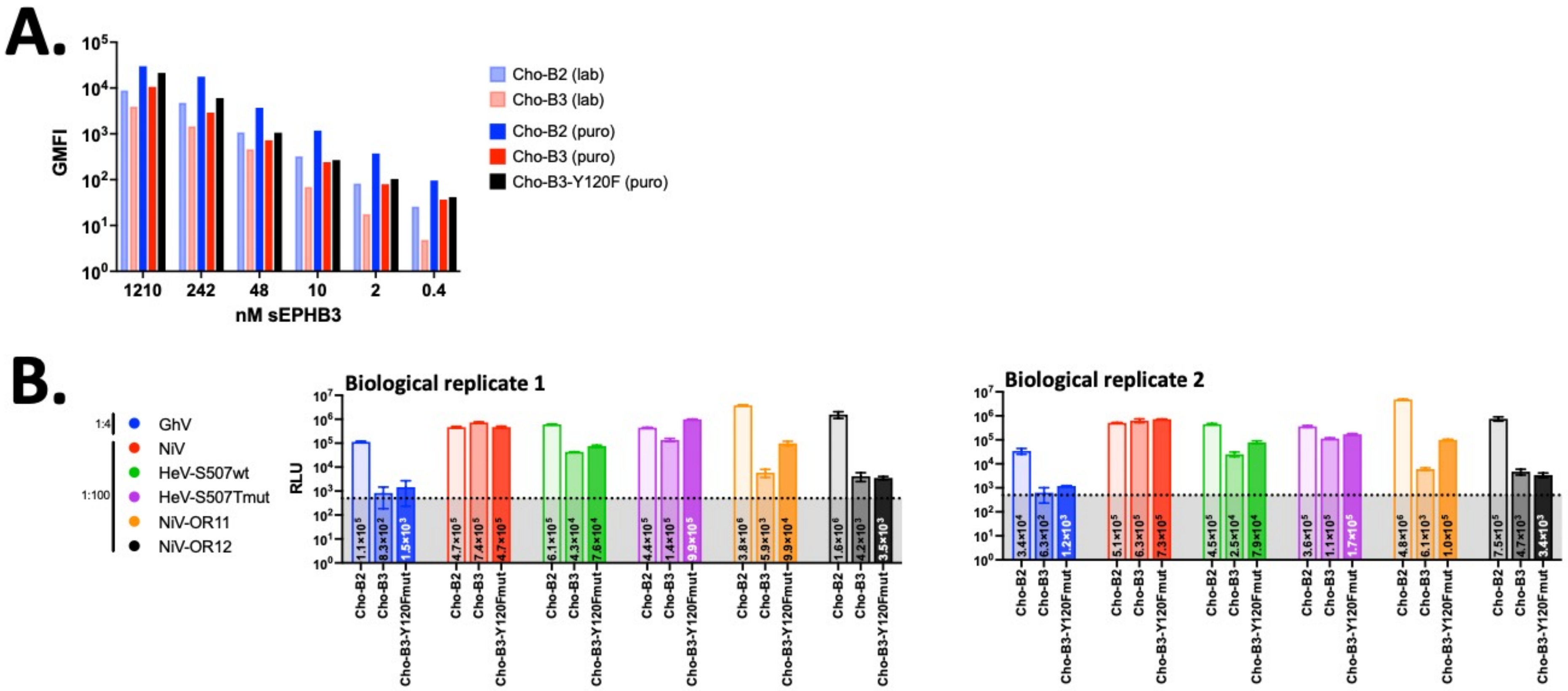

**Supplemental Figure 9. Use of and validation of CellProfiler for the automated quantification of HNV syncytia formation**

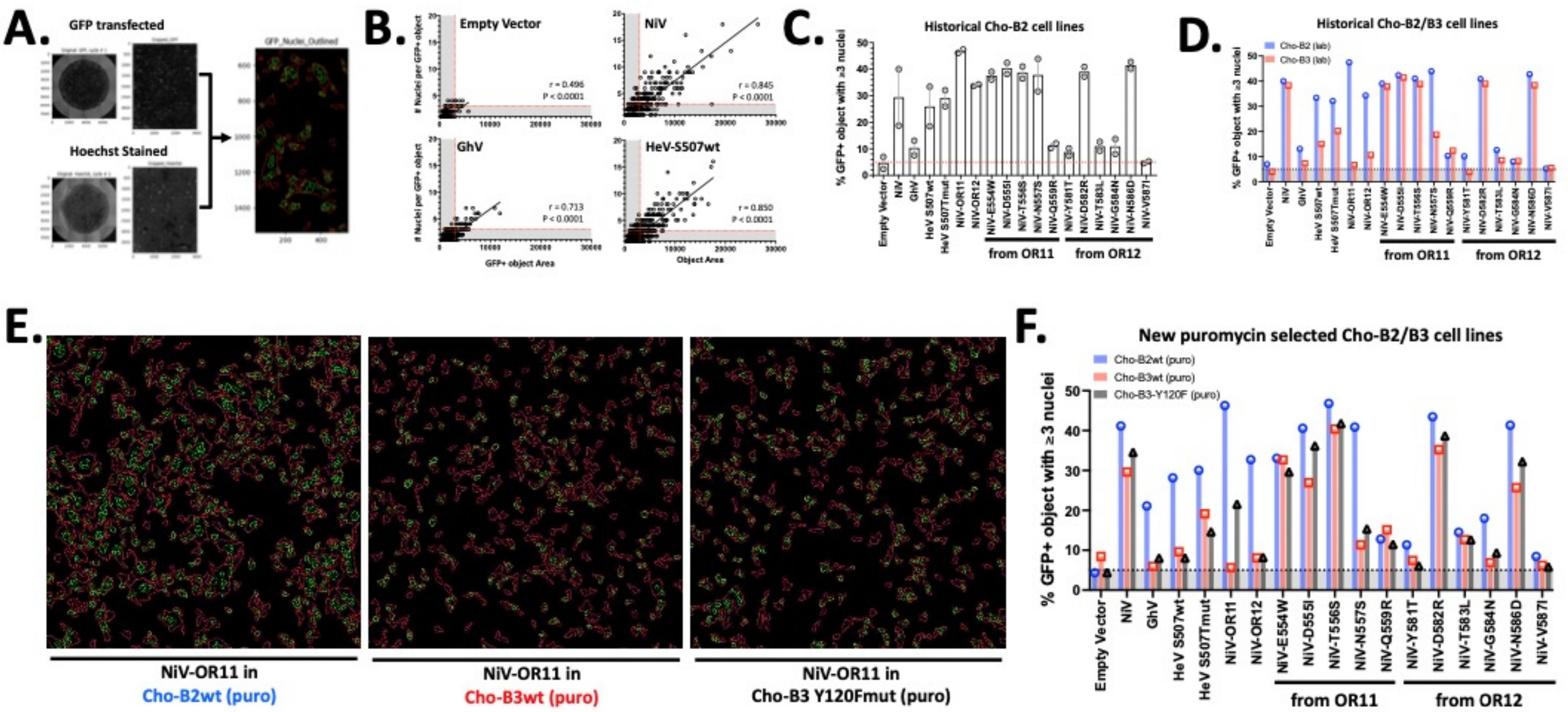

**Supplemental Figure 10. Recombinant NiV with selected mutants display decreased EFNB3 usage in vitro.**

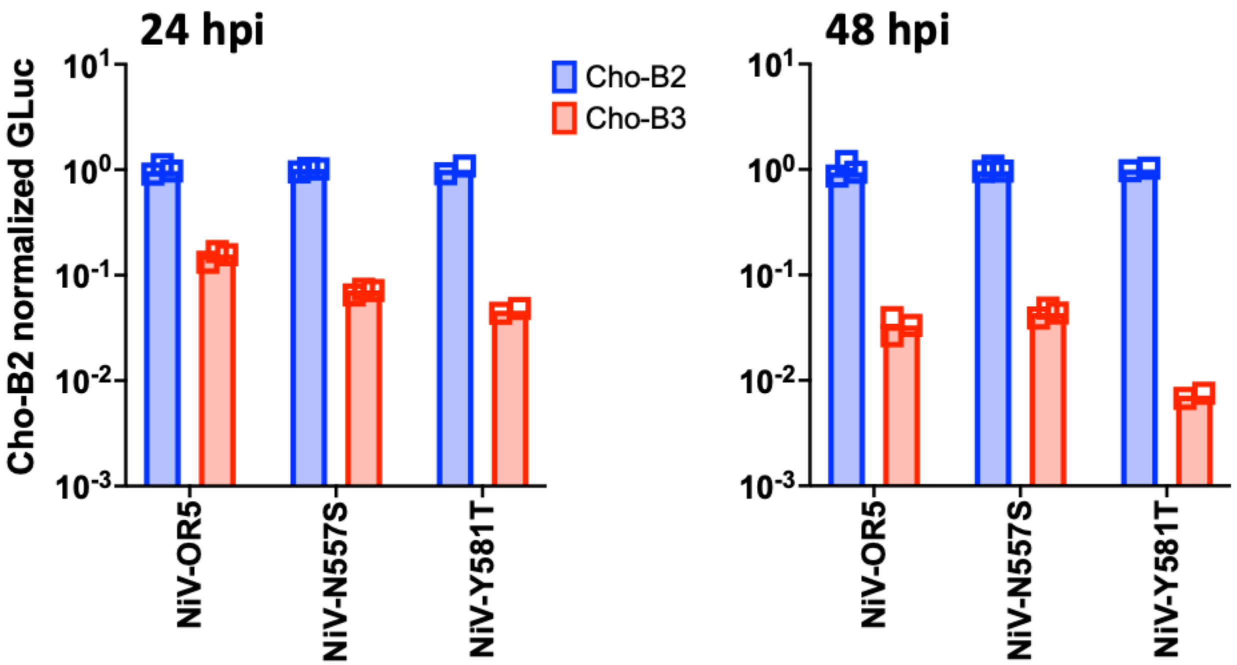

### Supplemental Table 1. Titers of HNVpp bearing NiV-OR mutants and point mutations

| | Construct | Avg. Cho-B2 Titer | Avg. Cho-B3 Titer | ChoB3 Titer/ChoB2 Titer |
|---|---|---|---|---|
| **Controls** | BALD | 4.47E+00 | 4.47E+00 | 1.0000 |
| | NiV | 6.91E+06 | 6.91E+06 | 1.0000 |
| | GhV | 1.34E+03 | 4.47E+00 | **0.0033** |
| | HeV-S507wt | 4.16E+06 | 2.82E+05 | 0.0678 |
| | HeV-S507Tmut | 1.41E+06 | 1.41E+06 | 1.0000 |
| **OR mutants** | NiV-OR1 | 6.91E+06 | 2.82E+05 | 0.0408 |
| | NiV-OR2 | 1.12E+04 | 4.69E+01 | 0.0042 |
| | NiV-OR3 | 6.91E+06 | 3.46E+06 | 0.5000 |
| | NiV-OR4 | 4.16E+06 | 8.46E+05 | 0.2034 |
| | NiV-OR5 | 6.91E+06 | 1.69E+05 | 0.0245 |
| | NiV-OR6 | 4.16E+06 | 8.46E+05 | 0.2034 |
| | NiV-OR7 | 6.91E+06 | 2.82E+05 | 0.0408 |
| | NiV-OR8 | 4.16E+06 | 2.82E+05 | 0.0678 |
| | NiV-OR9 | 6.91E+06 | 8.46E+05 | 0.1224 |
| | NiV-OR10 | 4.16E+06 | 8.46E+05 | 0.2034 |
| | NiV-OR11 | 6.91E+06 | 6.72E+03 | **0.0010** |
| | NiV-OR12 | 4.16E+06 | 1.34E+03 | **0.0003** |
| **OR11 point mutants** | NiV-E554W | 6.91E+06 | 6.91E+06 | 1.0000 |
| | NiV-D555I | 4.16E+06 | 4.16E+06 | 1.0000 |
| | NiV-T556S | 4.16E+06 | 6.91E+06 | 1.6611 |
| | NiV-N557S | 6.91E+06 | 1.12E+04 | **0.0016** |
| | NiV-Q559R | 3.37E+04 | 1.69E+05 | 5.0178 |
| **OR12 point mutants** | NiV-Y581T | 1.12E+04 | 4.47E+00 | **0.0004** |
| | NiV-D582R | 4.16E+06 | 1.41E+06 | 0.3389 |
| | NiV-T583L | 5.62E+04 | 2.82E+05 | 5.0178 |
| | NiV-G584N | 5.62E+04 | 3.37E+04 | 0.5996 |
| | NiV-N586D | 6.91E+06 | 4.16E+06 | 0.6020 |
| | NiV-V587I | 4.46E+02 | 2.24E+03 | 5.0224 |
